# Remote Detection of Red-Edge Spectral Characteristics in Floating Aquatic Vegetation

**DOI:** 10.1101/2024.12.24.630209

**Authors:** Aoi Murakami, Yu Komatsu, Kenji Takizawa

## Abstract

The vegetation red-edge of terrestrial plants is a key biosignature for detecting Earth-like habitable exoplanets. Although water is essential for plants, an excess of water can limit the distribution of terrestrial vegetation. On planets with extensive water coverage and limited land, floating vegetation on the water’s surface could serve as a crucial indicator of life. This study examined the spectral reflectance of floating plants across various scales, from individual leaves to lake-wide vegetation coverage. Our comparisons between individual leaves revealed that the red-edge of floating plants was equivalent to or even more pronounced than that of terrestrial plants. Although water can reduce plant reflectance, the naturally low reflectance of water enhances the detection sensitivity for floating vegetation. Seasonal changes, such as the proliferation of floating plants in summer and their decline in winter, lead to significant variations in lake reflectance. By analyzing satellite images of lakes and marshes over a five-year period, we confirmed that these seasonal variations in reflectance reliably indicate the presence of floating vegetation. The seasonal signal showed robustness to the effects of clouds, which pose another challenge on water-rich planets. We propose that floating vegetation be considered alongside, or even in place of, terrestrial vegetation in the search for extraterrestrial life.

## 1. Introduction

After a decade of intensive exoplanet surveys by the Kepler and TESS space telescopes, it has become evident that planets are ubiquitous in the universe. Some exoplanets are located at optimal distances from their central stars to sustain liquid water on their surfaces, classifying them as habitable planets (Seager, 2013). Ongoing follow-up observations by the latest space telescope, James Webb Space Telescope (JWST), and ground-based telescopes, are clarifying the planetary environments and their habitability. In the next decade, we anticipate achieving direct imaging of habitable planets (Fitzgerald et al., 2022), potentially enabling the detection of signs of life for the first time. Currently, the primary targets of biosignature surveys are habitable planets around M-dwarfs, smaller stars with lower temperatures than the Sun, and the most abundant stellar type in the universe (Scalo et al., 2007). Observing exoplanets around M-dwarfs provides valuable insights into planetary environments through transit spectroscopy, enabling the characterization of atmospheric composition. The understanding of planetary environments and the advancement in techniques, such as adaptive optics (Males et al., 2014), lead to future direct imaging of surface biosignatures around Sun-like (F, G, K) stars, which offer larger planet-star separations and more stable stellar fluxes for their habitable planets compared to M-dwarfs (O’Malley-James & Kaltenegger, 2018a).

If Earth is considered a representative habitable planet, molecular oxygen produced by green plants is a prominent gaseous biosignature detectable remotely. However, the risk exists of a false positive for life, as oxygen can accumulate in the atmosphere from the abiotic oxidation of water (Luger and Barnes, 2015, Meadows et al., 2018, Narita et al., 2015). To confirm the presence of plant-like life, additional biosignatures are necessary. Plant leaves absorb visible light via photosynthetic pigments (Sims and Gamon, 2002) and strongly reflect near-infrared (NIR) radiation through internal leaf scattering (Knipling, 1970). This results in a reflection gap between red light and NIR (around 700 nm), known as the vegetation red-edge (VRE). Satellite remote sensing of terrestrial vegetation can track the VRE, enabling the assessment of vegetation distribution and physiological state (Rouse et al., 1974). Similar methods could be applied in the direct imaging of exoplanets in the near future, with the VRE expected to serve as a surface biosignature (Kiang et al., 2007; Knipling, 1970; Montañés-Rodríguez et al., 2006; Seager et al., 2005).

Plants or phototrophs are likely to exist on habitable exoplanets, as they can grow autotrophically in the presence of water and light. While the optical properties of gaseous biosignature molecules like oxygen are physically determined, the spectral reflectance of phototrophs may vary biologically depending on their growth environment. Around M-dwarfs, with their extreme light conditions, the VRE might shift to longer wavelengths compared to that around the Sun (Tinetti et al., 2006). However, according to findings from our research group, an Earth-like VRE could develop through the evolution of aquatic phototrophs into land vegetation (Takizawa et al., 2017). Further studies are needed to determine the exact wavelength position of the red-edge around M-dwarfs. Nevertheless, a wide observational wavelength range from the visible to the near-infrared should facilitate detection despite any variations.

One practical challenge in observing the VRE on exoplanets is the limited distribution of plants. Earthshine observations by Seager et al. (2005) suggest that VRE observability is significantly influenced by the surface area covered by vegetation. On exoplanets, insufficient continental landmasses could hinder VRE observations. If water presence is limited on habitable planets around M-type stars (Tian and Ida, 2015; Wandel, 2018), the absence of large continents might not be an issue, but such planets are less likely to support life. Conversely, rocky planets around M-dwarfs can maintain a moderate amount of water over extended periods if water is produced in their primordial atmospheres (Kimura and Ikoma, 2022). While the likelihood of life increases with the availability of water, water-rich environments may pose challenges to VRE detectability. The recent discovery of water vapor in the terrestrial planet-forming zone by JWST supports the latter scenario (Perotti et al., 2023).

On habitable exoplanets with ample liquid water, land vegetation may be scarce even if a thriving oceanic biosphere exists. Submerged aquatic plants and algae are prevalent in both marine and freshwater environments on Earth, but their remote detection is highly constrained due to light absorption by the water column (Cho et al., 2008; Han and Rundquist, 2003). Only floating plants and emergent plants with floating leaves are easily identifiable via satellites. Although their distribution is currently limited to freshwater regions near land on Earth, global blooms of floating macrophytes may have occurred in the Eocene Arctic Ocean, known as Azolla events (Brinkhuis et al., 2006; Neville et al., 2019). The reflectance spectra of floating leaves are similar to those of terrestrial plants, but the background reflectance of water differs significantly from that of soils. Consequently, the observed reflectance spectrum of floating vegetation varies considerably based on water coverage. In Earth observations using satellites, this variability is advantageous for classifying surface regions through time series analysis (Hou et al., 2018; Wang et al., 2023).

In this paper, we evaluate the light reflectance properties of various types of floating plants at different observational scales and explore the potential for extending remote VRE detection to exoplanets. For the purposes of this study, “floating plants” refer to aquatic plants that spread their leaves above the water’s surface. We first measure a variety of floating plants under different conditions to identify unique reflectance characteristics distinct from those of terrestrial and submerged vegetation. Based on these characteristics, we assess whether floating plants can be detected from the reflectance spectra of entire lakes. By integrating satellite remote sensing data with topographic and vegetation data to obtain lake reflectance data, and employing unsupervised machine learning techniques, we analyze annual variation patterns that characterize floating vegetation and evaluate the impact of cloud cover. While previous studies in aquatic plant ecology and remote sensing have aimed to enhance methods for species classification and biomass quantification, our study focuses on the common light reflection properties of floating vegetation and the robustness of these signals for potential future detection on habitable exoplanets.

## 2. Materials and Methods

We investigated the light reflection characteristics of floating plants at different scales, ranging from individual leaves (cm^2^) to natural vegetation areas (km^2^). Small-scale, precise spectral measurements were conducted using fiber spectroscopy and a hyperspectral camera in the laboratory. Medium-scale measurements of various floating plants under sunlight were performed with a multi-band sensor designed for vegetation surveys. Large-scale and long-term analyses were carried out using satellite data from multispectral imagers.

### 2.1 Plant materials for reflection spectral measurements

For the laboratory measurements of reflectance spectra, we used seven floating plants (*Lemna minor*, *Azolla filiculoides*, *Salvinia molesta*, *Limnobium laevigatum*, *Eichhornia crassipes*, *Nymphaea spp*., and *Trapa japonica*), one submerged aquatic plant (*Egeria densa*), one green alga (*Chlamydomonas reinhardtii*), and four terrestrial plants (*Arabidopsis thaliana*, *Ligustrum lucidum*, *Verbascum thapsiforme*, and *Kalanchoe CLONEKOE*). The green alga was cultivated in a Tris-acetate-phosphate medium at 23℃ under illumination of 40 μmol photons m⁻^2^s⁻¹ with rotary shaking. The plants were grown in fertilized water or soil in a greenhouse at 20℃–30℃, except for the larger plants (*Nymphaea spp*., *Trapa jeholensis*, and *Ligustrum lucidum*), whose leaves were collected from specimens growing in the field around our institute. Plants pictures are shown in Figure S1.

The reflectance spectrum of natural floating vegetation was measured at two locations near our institute: Oroike (34.96 N, 137.20 E) and Lake Suwa (36.06 N, 138.10 E). Oroike is a relatively small (0.01 km^2^, depth < 2 m) warm pond with an average winter temperature of about 5℃. Horticultural varieties of hardy water lilies (*Nymphaea spp.*) form colonies in shallow waters (less than 1 m). Although water lilies are classified as emergent plants with rhizome networks at the lake bottom, they are treated here as floating plants for convenience. Lake Suwa is a larger (13.3 km^2^, maximum depth < 7 m) colder pond with an average winter temperature of −1℃. Water chestnut (*Trapa japonica*) forms large colonies, approximately 1 km long, along the coastal areas at water depths less than 3 m.

### 2.2 Quantification of chlorophyll concentration

Following leaf-reflectance measurements, leaf thickness, fresh weight, and chlorophyll concentration were measured for each sample. For larger plants, leaves were separated, while for smaller plants, the entire plant body was used for chlorophyll measurements. A small amount of leaf tissue (0.02–0.2 g) was frozen in liquid nitrogen and ground with 3–10 mL of 80% acetone. Total chlorophyll (a + b) concentration was measured spectroscopically using the equation of Porra et al. (1989).

### 2.3 Spectral reflectance measurement using a fiber optic spectrometer

Vertical reflectance of individual leaves was measured using a duplex fiber reflection probe (RP28; Thorlabs) connected to a tungsten halogen light source (HL-2000-HP; Ocean Optics) and a Charge-Coupled Device (CCD) array spectrometer (FLAME-S-VIS-NIR-ES; Ocean Optics). The sample end of the Y-bundle was placed 15 mm vertically above the leaf surface. The fiber optic spectrometer detects reflected light within the 350 to 1000 nm range, with a resolution of 1.5 nm. Spectral reflectance was calculated by comparing the sample spectrum with a reference spectrum from a standard (USRS-99-010; Labsphere). Detached leaves were placed on a sample stage, covered with black paper containing a 1 cm diameter hole positioned beneath the fiber probe, to measure the reflectance of the exposed area. The sample stage height was adjusted to keep the detection distance constant among the different leaves. Each measurement was repeated three times per leaf, with the leaf rotated 120 degrees on the sample stage, and the results were averaged.

### 2.4 Spectral imaging of plants using a hyperspectral camera

Small-scale spectral reflectance of aquatic plants was measured using a hyperspectral camera (KD-1; EBA Japan Co., Ltd.). The Complementary Metal Oxide Semiconductor (CMOS) sensor recorded hyperspectral images in 101 bands between 400 and 900 nm, as described by Kohzuma et al. (2021). Uniform incident light in the 400–800 nm range was provided by two xenon lamps (XC-100B; Seric Ltd.), placed on either side of the sample at a 45° angle. A plastic cup lined with black paper was filled with water to a depth of 3 cm, and the floating plants were placed on the water (see Figure 1A). The camera was set up 30 cm vertically above the water surface and scanned an area of 9 cm x 9 cm (Figure S2). Raw images of reflected radiation were converted to reflectance images using a diffuse reflectance standard (USRS-99-010; Labsphere). The reflectance from a central area of 3.6 cm x 3.6 cm (25,600 pixels) was averaged. Each measurement was repeated three times per sample, with the sample rotated 120 degrees on the sample stage, and the results were averaged.

**FIG. 1.**
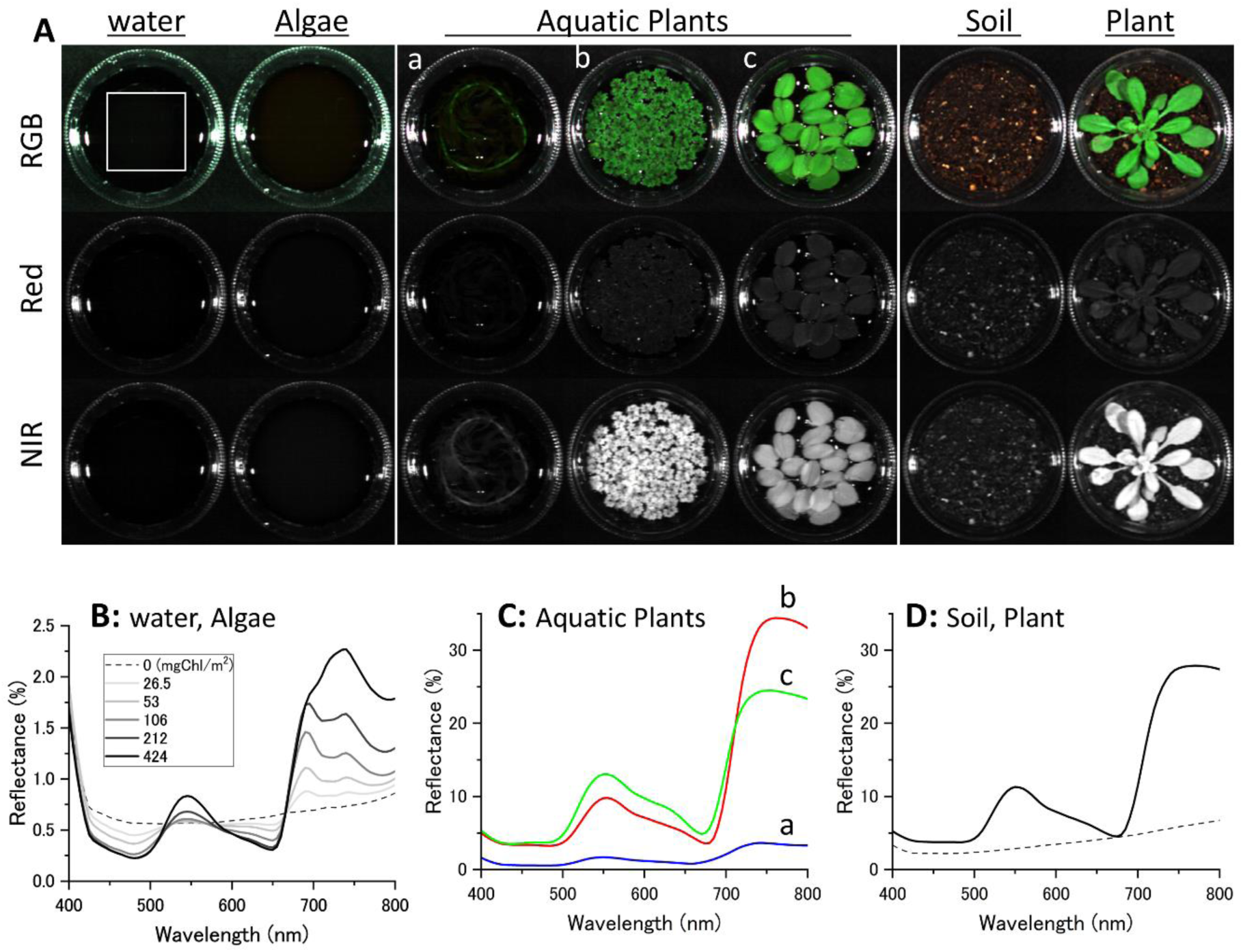
Spectral image and reflectance spectrum of algae, aquatic plants, and a terrestrial plant. **A.** Spectral images were captured by a hyperspectral camera. Color images (RGB) and reflectance images at 670 nm (Red) and 800 nm (NIR) for each plant are shown. The labels a, b, and c on the images of aquatic plants correspond to the symbols in the reflectance spectra. The upper left square (3.6 x 3.6 cm) represents the measurement area of the reflectance in each image. **B.** Reflectance spectra of green algae (*Chlamydomonas reinhardtii*). The dashed line represents the reflectance of water. Solid lines with different densities represent the reflectance spectra of green algae with varying cell densities. Chlorophyll concentrations were 27, 53, 106, 212, and 424 mg/m^2^. **C.** Reflectance spectra of aquatic plants. Graphs a (blue line), b (green line), and c (red line) represent *Egeria densa* (223 mg Chl/m^2^), *Azolla filiculoides* (228 mg Chl/m^2^), and *Salvinia molesta* (249 mg Chl/m^2^), respectively. **D.** Reflectance spectra of a terrestrial plant (*Arabidopsis thaliana*, 132 mg Chl/m^2^). The dashed line represents the moist soil used for growing the plant.

Under the same conditions, spectral reflectance images of green algae (*Chlamydomonas reinhardtii*) and a submerged plant (*Egeria densa*) were measured. The density of the submerged plant, converted based on chlorophyll concentration per area, was set to be about the same as that of the floating plants. For the green algae, measurements were conducted across a wide range of concentrations to examine concentration-dependent changes in reflectance. For comparison with a terrestrial plant, *Arabidopsis thaliana* was planted in the plastic cup filled with soil instead of water and measured during the rosette growing stage, ensuring that all leaves were spread on the soil surface.

### 2.5 Spectral imaging of floating plant populations

Spectral reflectance patterns of floating vegetation under sunlight were measured using a multispectral camera (RedEdge-M; MicaSense). The camera’s five CMOS sensors simultaneously captured upwelling radiance from the plants at 475, 560, 668, 717, and 840 nm. Ambient spectral irradiance was measured concurrently by an attached downwelling light sensor. Relative reflectance, calculated from the measurements of the two sensors, was converted to absolute reflectance using a calibrated reflectance panel (MicaSense).

Spectral reflectance of various plant populations was measured for plants floating in a water tray (85 x 55 cm) with a water depth of 15 cm. The black plastic tray was lined with black cloth to minimize bottom reflection. The camera placed 85 cm vertically above the water surface had a field of view of 74 cm x 55 cm, which covered 88% of the water tray (Figure S3).

The five-band images were aligned and processed using a Python script provided by MicaSense (https://micasense.github.io/imageprocessing/Alignment.html). Since aligning the five-wavelength images at close range is challenging, each surface area was captured three times, with the floating plant positions altered between captures, and the results were averaged. A surface area of 807 cm^2^ (240,000 pixels) was selected for reflectance analysis, avoiding areas with sun glitter and camera shadows.

Spectral reflectance of floating vegetation in natural habitats was measured using the camera mounted on a quadcopter drone (Matrice 100; DJI). We studied the two lakes indicated above over two years (2021-2023). All aerial photographs were taken under clear skies between 11:00 AM and 1:00 PM. While flying at an altitude of 100 m, the camera captured a series of nadir shots with 75% overlap in both the direction of flight and the lateral direction. Sequential radiance images were converted into orthomosaics and reflectance maps using image processing software (Pix4D mapper; Pix4D). Spectral reflectance and vegetation indices in selected areas were calculated using a geographic information system (QGIS 3.0).

### 2.6 Satellite remote sensing dataset for analyzing lake-wide aquatic vegetation

Long-term reflectance spectroscopy patterns of lakes and marshes throughout Japan were analyzed using satellite remote sensing. The data for remote sensing were obtained using the MSI sensor on Sentinel-2, part of the Copernicus program operated by the European Space Agency (ESA). Sentinel-2, an optical satellite in a sun-synchronous orbit, continuously collects observation data primarily over land between 56°S and 84°N latitude. Sentinel-2A (S2A) was launched in 2015, and Sentinel-2B (S2B) has been in operation since 2017, each with a 10-day revisit period. The MSI sensor detects wavelengths from visible light to infrared across 13 bands (https://sentiwiki.copernicus.eu/web/s2-mission). The analysis mainly utilized the red light of Band 4 (S2A: 664.6 nm, S2B: 664.9 nm) and the near-infrared of Band 8 (S2A: 832.8 nm, S2B: 832.9 nm), both with a spatial resolution of 10 m. While the Sentinel-2 bands in the region closer to the peak NIR reflectance of plants (750-760 nm), namely B5 at 705 to B7 at 783 nm, are more suitable for sensitive vegetation detection, their lower spatial resolution of 20 m limits their effectiveness for detecting floating vegetation in smaller-scale environments such as lakes and marshes. Consequently, Bands 4 and 8 were selected for vegetation detection in this analysis. The central wavelengths of these bands closely match those of the Multiband camera described in the previous section, allowing a direct comparison between the satellite data analysis and the results from our laboratory and field measurements.

Sentinel-2 provides two datasets with different levels of correction: Level-1C (L1C) and Level-2A (L2A). Further details of each product are described in Section 2.8. Data from 2019 to 2023 were used, as these years showed relatively stable quality for both L1C and L2A products. The datasets were accessed via Google Earth Engine (GEE), a cloud-based geospatial processing platform (Gorelick et al., 2017). Both raw and cloud-masked data from L1C and L2A were utilized in the analysis. The cloud masking process employed the “Cloud Score + S2_HARMONIZED V1” product provided by GEE, with a threshold of 0.6 applied to the “cs_cdf” (Cloud Score Cumulative Distribution Function) band, effectively removing cloud-contaminated pixels from the analysis (Pasquarella et al., 2023). The Normalized Difference Vegetation Index (NDVI) was calculated using Sentinel-2 MSI bands 4 and 8. The study focused on 148 sites, selected by filtering geographic information for reported aquatic plant community sites (Kato et al., 2024) from the Ministry of Land, Infrastructure, Transport and Tourism’s lake and pond dataset (W09-05_GML; https://nlftp.mlit.go.jp/ksj/gml/datalist/KsjTmplt-W09-v2_2.html). These sites included both freshwater and brackish lakes, all known to host aquatic plant communities. In addition, when collecting images via GEE, the coordinate system was standardized to WGS 84. These 148 locations are listed in Table S1.

### 2.7. Time series analysis of NDVI

The seasonal variation in NDVI time series data over five years for 148 locations was reconstructed using the Harmonic Analysis of Time Series (HANTS) method (Malamiri et al., 2020; Roerink et al., 2000). Fast Fourier Transform (FFT) and power spectrum analysis were employed to determine the periodicity in each time series dataset, with these measurements conducted using L2A data. The analysis showed that 117 out of the 148 locations exhibited a distinct annual cycle. HANTS fitting was then applied to these 117 locations, using data extracted from different levels of datasets as described in the following section. Before applying HANTS fitting, average values were calculated for each data acquisition date, which were then integrated into a representative one-year time series dataset. To standardize the temporal resolution of the data, the following factors were considered: (1) the occurrence of leap years, (2) overlapping grids in large lakes, and (3) data acquisition at 5-day intervals by the two Sentinel-2 satellites. The one-year period was divided into 5-day intervals, with the Day of Year (DOY) defined as [5, 10, 15,…, 365], where DOY 5 represents the average data value acquired between January 1 and January 5. This preprocessing resulted in a temporally homogeneous one-year NDVI time series dataset, optimized for HANTS analysis. During the HANTS analysis, the “number of frequencies,” a critical parameter, was set to 4 (Zhou et al., 2015), selected to effectively capture annual vegetation dynamics while adequately avoiding overfitting. Additionally, the “fit error tolerance” parameter was set to 0.05 to balance fitting accuracy and robustness. This approach enabled the extraction of the main frequency components of seasonal variations, minimizing the influence of noise.

### 2.8 Unsupervised machine learning: clustering of lakes with floating vegetation

Following HANTS fitting, clustering analysis was performed using the K-shape method. Locations with consistently positive or negative NDVI values were excluded, as consistently positive values could indicate interference from terrestrial vegetation near the lake, while consistently negative values might suggest either the absence of vegetation or its presence at a very small scale. This filtering reduced the number of locations from 117 to 92. Next, among these 92 locations, those with a maximum NDVI value of 0.1 or higher, the threshold for sparse grassland, were selected based on the cloud-masked L2A data (Ya’acob et al., 2014). As a result, 72 locations were chosen for clustering analysis (Table S2).

For clustering, a dataset derived from L2A products with additional cloud masking applied (L2A-CM) was used to accurately capture the NDVI fluctuations of vegetation. K-shape is a clustering technique that focuses on the shape of time series data. It interprets time series data as waveform functions and assesses similarity based on a distance measure that considers the phase and amplitude of the waveforms. Due to these characteristics, K-shape can potentially outperform conventional K-means in time series clustering applications (Paparrizos and Gravano, 2015). The optimal number of clusters was determined using the elbow method, leading to the selection of three clusters (Figure S4). The K-shape clustering was implemented using the Python library “tslearn” (Tavenard et al., 2020).

### 2.9 Assessment of effects by cloud and atmosphere

This analysis utilized three hierarchical levels of Sentinel-2 datasets: L1C, L2A, and L2A-CM. L1C data, representing top-of-atmosphere reflectance, undergoes processing up to orthorectification (https://sentiwiki.copernicus.eu/web/s2-processing#S2Processing-L1CAlgorithmsS2-Processing-L1C-Algorithmstrue). L2A data is generated by applying atmospheric and topographic corrections to L1C data using the libRadtran radiative transfer model (https://step.esa.int/thirdparties/sen2cor/2.10.0/docs/S2-PDGS-MPC-L2A-ATBD-V2.10.0.pdf), providing bottom-of-atmosphere reflectance. L2A-CM data is derived from L2A by excluding cloud-contaminated pixels, as detailed in Section 2.6.

A comparative analysis of these datasets was conducted to quantify the effects of cloud and atmospheric attenuation caused by atmospheric molecules. The Cloud Probability band in L2A, with an annual average of approximately 40%, was used to characterize cloud cover.

Atmospheric gaseous effects were assessed by comparing L1C and L2A datasets, while overall attenuation was quantified by comparing L1C and L2A-CM, simulating conditions similar to astronomical observations.

For each dataset, the annual maximum NDVI (NDVI_Max_) and the difference between maximum and minimum NDVI (ΔNDVI) were calculated. Attenuation rates were evaluated by analyzing the frequency distribution and median values of NDVI_Max_ and ΔNDVI.

## 3. Results

### 3.1. Spectral reflectance of water-floating leaves

In the initial step of the multi-scale analysis, we compared the spectral reflectance properties of individual leaves from water-floating plants with those of terrestrial plants. Seven species of floating plants and four species of terrestrial plants, each with varying leaf morphologies, were selected for this comparison (see Materials and Methods and Figure S1). Light reflectance spectra of single-layered leaves were measured using a fiber spectrometer (FLAME-S-VIS-NIR-ES; Ocean Optics), as shown in Figure S5. The characteristics of the diverse leaves are summarized in Table S2. All leaves exhibited distinctive red-edge features, characterized by low reflectance in the red light and high reflectance in the near-infrared radiation (NIR). The red-edges of the floating plants were found to be as pronounced as, or even more pronounced than, those of the terrestrial plants. The NIR reflectance of vascular plant leaves varies significantly depending on leaf morphology (Slaton et al., 2001) and is influenced by factors such as leaf thickness, the cuticular layer, and trichome density. The floating plant leaves used in this study also displayed these morphological characteristics, and high NIR reflectance was observed. The wavelength positions of the red-edge, determined from the peak in the first derivative of the reflectance spectrum, showed minimal variation with leaf morphology and were consistently within the range of 707–715 nm.

However, these results do not represent reflectance in habitats where floating leaves are wet. Therefore, we measured the reflectance of floating plants (*Azolla filiculoides* and *Salvinia molesta*) on the water surface and compared them to a green alga (*Chlamydomonas reinhardtii*) culture, a submerged aquatic plant (*Egeria densa*) in water, and a terrestrial plant (*Arabidopsis thaliana*) on soil to determine the effect of the water background. Spectral images were captured using a hyperspectral camera (KD-1; EBA Japan Co., Ltd.), as shown in Figure 1A. Due to NIR absorption by water, the NIR reflectance of the unicellular green alga was minimal. Although the red-edge features of algal water increased with cell density (Figure 1B), they remained an order of magnitude smaller compared to the terrestrial plant (Figure 1D). The red-edge position of the algae was in the range of 668–680 nm, slightly shorter compared to the plant leaves. The submerged aquatic plants consist of distinct leaf and stem structures, but only the stem was evident in the RGB and NIR images in Figure 1Aa. Submerged leaves were obscured when imaging sensitivity and display scale were optimized for interspecies comparison with floating plants (Figure S2). The red-edge feature of the submerged aquatic plant (Figure 1Ca) was more significant than the green algae but still did not reach the levels observed in the terrestrial plant. Floating plants (Figure 1Cb,c) on the water surface displayed a pronounced red-edge comparable to that of the terrestrial plant. Unlike the results from the reflectance spectral measurements of dry leaves, the NIR reflectance of *Salvinia* (Figure 1Cc) was lower than that of *Azolla* (Figure 1Cb) and the terrestrial plant. Leaves of the terrestrial plant measured on soil were dry on both sides, whereas the leaves of floating plants measured above water were wet on the underside. Given its larger leaves, *Salvinia* was likely more affected by wetting. The red-edge position of the floating plants was in the range of 705–710 nm, which is equivalent to that of the terrestrial plant.

### 3.2 Spectral reflectance measurements of floating plants population

In the next step, the light reflection properties of floating plant populations were investigated in detail. Five different types of floating plants—*Lemna minor*, *Azolla filiculoides*, *Salvinia molesta*, *Limnobium laevigatum*, and *Eichhornia crassipes*—were floated on a water tray (85 x 55 cm) at varying densities, and spectral reflectance images were taken under sunlight using a 5-band image sensor (RedEdge-M, MicaSense) as described in the Materials and Methods and Figure S3. After capturing the reflectance images, fresh weight (FW) and chlorophyll concentration ([Chl]) of the plants were measured.

The coverage ratio of the water surface by floating plants was determined from the percentage of pixels with NIR reflectance higher than a species-specific threshold. Small plants (*Lemna* and *Azolla*) have thalli and roots a few millimeters long spread as a thin layer over the water surface. The relatively larger plants (*Limnobium* and *Eichhornia*), with leaves, stems, and root structures a few centimeters long, protruded the upper surface or both surfaces of their leaves above the water, forming clumps. When leaf coverage was plotted against FW, smaller plants tended to cover larger areas with less mass compared to larger plants (Figure 2A).

**FIG. 2.**
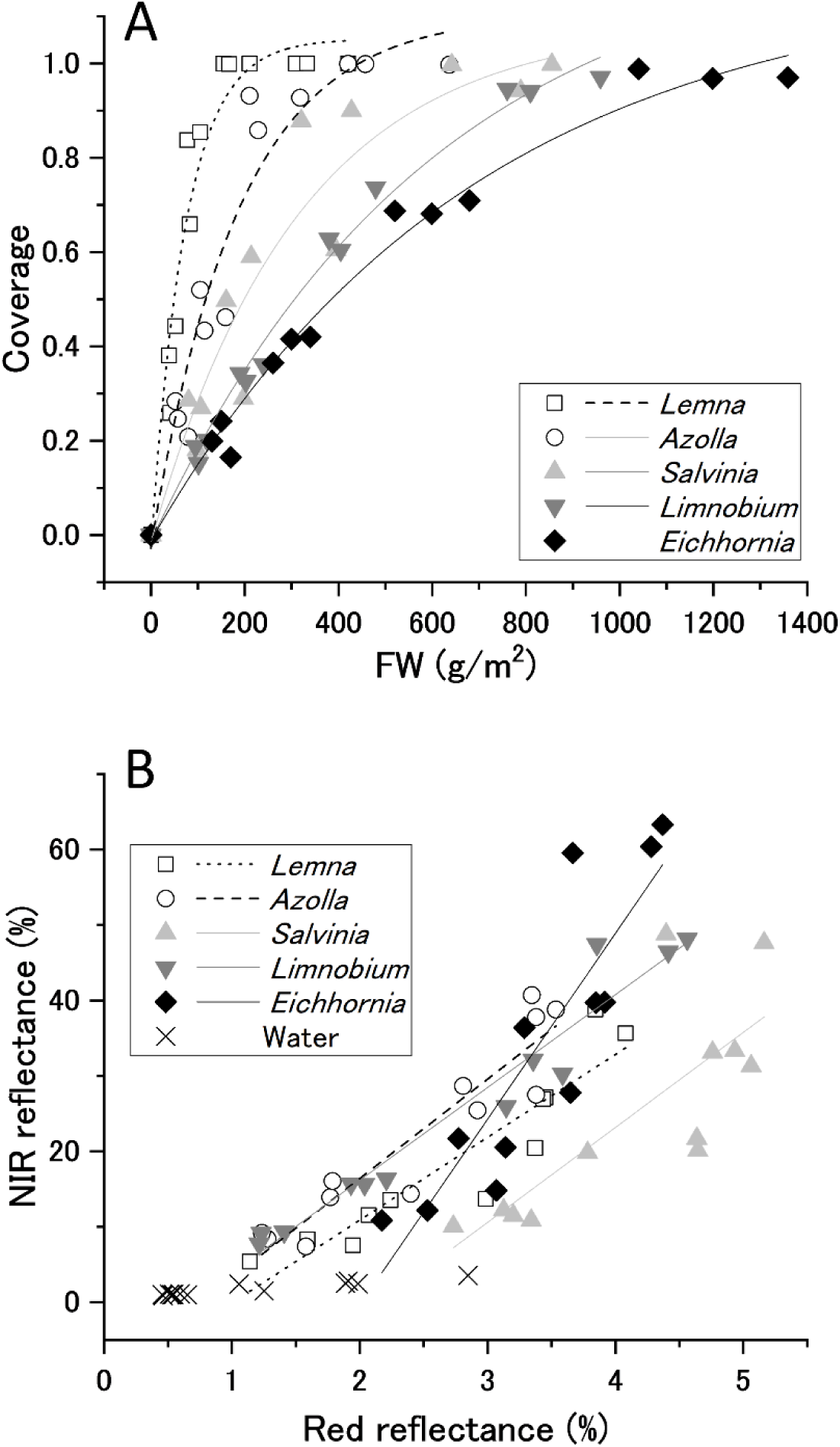
Diffusion patterns and spectral reflectance properties of floating plants of different types and densities. Different symbols represent different species: *Lemna minor* (□), *Azolla filiculoides* (○), *Salvinia molesta* (▴), *Limnobium laevigatum* (▾), *Eichhornia crassipes* (◆), *and water* (╳). **A.** Percent coverage of the water surface by different floating plants plotted against fresh weight and fitted by the Yield-fertilizer model in agriculture (y = a + br^x^). **B.** NIR reflectance plotted against red reflectance, with a linear fit applied.

The reflectance spectra of floating plant populations are shown in Figure S6. Under conditions of dense surface coverage by the plants, all species exhibited pronounced red-edge features. As plant density decreased, reflectance decreased across all bands and for all species. The reflectance of the empty water surface was lower than that of the plants at all wavelengths. When NIR (840 nm) reflectance was plotted against red (668 nm) reflectance, a monotonically increasing relationship was observed, with NIR reflectance being an order of magnitude greater than red reflectance (Figure 2B). Although the increase in red reflectance due to the floating plant cover is minor, it is a unique feature not seen in other vegetation. Terrestrial vegetation or aquatic algae decreases red reflectance compared to soil or water surface (Figure 1B).

To quantitatively assess changes in red-edge size due to variations in vegetation density on the water surface, we compared three indices commonly used to assess terrestrial vegetation: the ratio vegetation index (RVI) (Jordan, 1969), the difference vegetation index (DVI) (Tucker, 1979), and the normalized difference vegetation index (NDVI) (Rouse et al., 1974). These indices are calculated from the reflectance in the red *(R*_red_) and NIR (*R*_NIR_) bands according to the following equations:

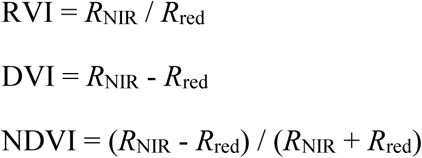

The indices for five species at different densities were calculated and plotted against FW, [Chl], and surface coverage ratio (Figure 3).

**FIG. 3.**
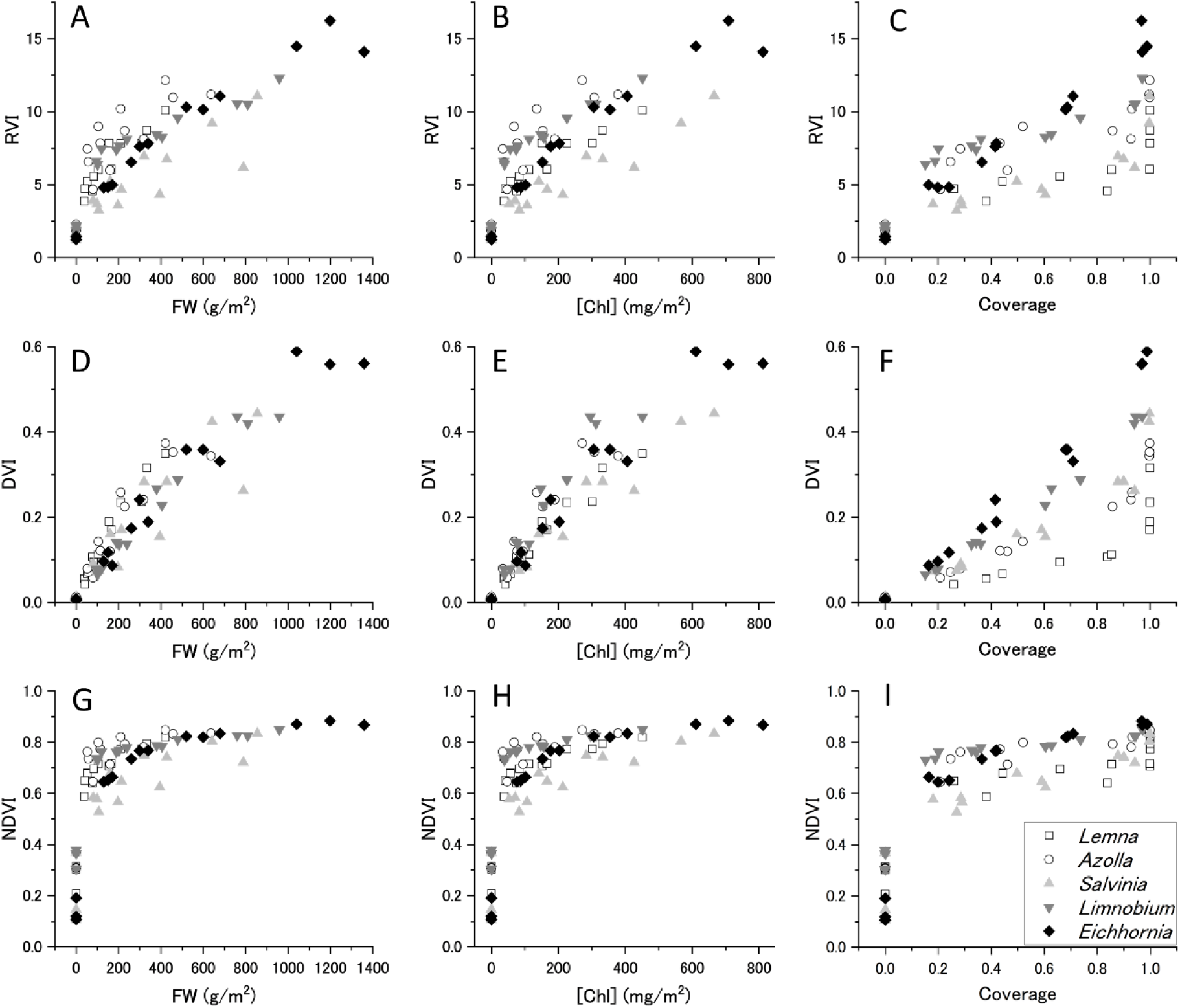
Vegetation indices of floating plant populations. RVI (A, B, and C), DVI (D, E, and F), and NDVI (G, H, and I) are plotted against fresh weight (FW), chlorophyll concentration ([Chl]), and plant coverage of the water surface (Coverage). Symbols represent different species, as shown in Figure 2.

RVI was roughly proportional to FW, [Chl], and coverage, with both inter- and intra-species variation (Figure 3A, B, C). For common terrestrial vegetation, RVI is highly sensitive at relatively high plant coverage, as *R*_red_ decreases with increasing *R*_NIR_. In contrast, for floating plants, both *R*_red_ and *R*_NIR_ increased with increasing coverage, indicating high detection sensitivity at low plant coverage and rapid saturation with increasing coverage.

DVI was also proportional to FW and [Chl], with less inter- and intra-species variability than RVI (Figure 3D, E). DVI exhibited high linearity with plant coverage, except for small plants at maximum density (Figure 3F). Larger plants showed higher DVI values relative to coverage. The smallest plant (*Lemna*) exhibited high DVI when the plants overlapped above the surface saturation density.

Among the three indices, NDVI was the most sensitive to small FW, [Chl], and coverage (Figure 3G, H, I). Previous research has reported that NDVI shows large values that deviate significantly from linearity with plant coverage when mixed pixel calculations include the bare water area (Chen, 1999; Price, 1992). In this study, we calculated NDVI from the red and NIR reflectance averaged over the entire measurement area (240,000 pixels) rather than each pixel, confirming high NDVI values (∼0.6) even at the lowest water coverage (∼20%) for all species.

These results indicate that DVI is suitable for estimating the amount and coverage of known floating plants. Conversely, RVI and NDVI have higher detection sensitivity when the vegetation cover ratio is low. In particular, NDVI may be able to detect aquatic vegetation with less than 20% coverage, regardless of the morphology of the floating plants. NDVI is thus the best index for detecting floating plants when the water surface coverage is low.

### 3.3 Spectral reflectance measurements of natural floating vegetation

In the third step, we studied the light reflectance characteristics of two lakes in both summer and winter. The five-band reflectance images of water areas covered by floating vegetation were compared with those of empty water and the terrestrial vegetation surrounding the lakes. Information about the survey sites is provided in the Materials and Methods section and Figure S7.

Figures 4A and 4B show the reflectance spectra of a small reservoir and the surrounding forest. During the summer (Figure 4A), hardy water lilies (*Nymphaea spp.*) thrived in the shallow (<1 m) peripheral areas (Figure S7A). The NIR reflectance of water lilies was as high as that of the mixed evergreen and deciduous broad-leaved forest surrounding the lake. The visible light reflectance of the waxy floating leaves was slightly higher than that of the tree leaves. The red-edge of the water lily community was evident in contrast to the bare water surface but was slightly less pronounced than that of the forest. In winter (Figure 4B), the water lilies disappeared (Figure S7B), revealing a low proportion of submerged plants. Although the riparian forest shed some leaves during winter, it retained red-edge features. The water surface reflectance remained low across all wavelengths throughout the season. The NIR reflectance of water without floating leaves was slightly higher than the red reflectance. with this difference being more pronounced in summer. Besides water lilies, perennial aquatic plants (*Cabomba caroliniana*) were observed throughout the pond, contributable to the tiny red-edge features.

**FIG. 4.**
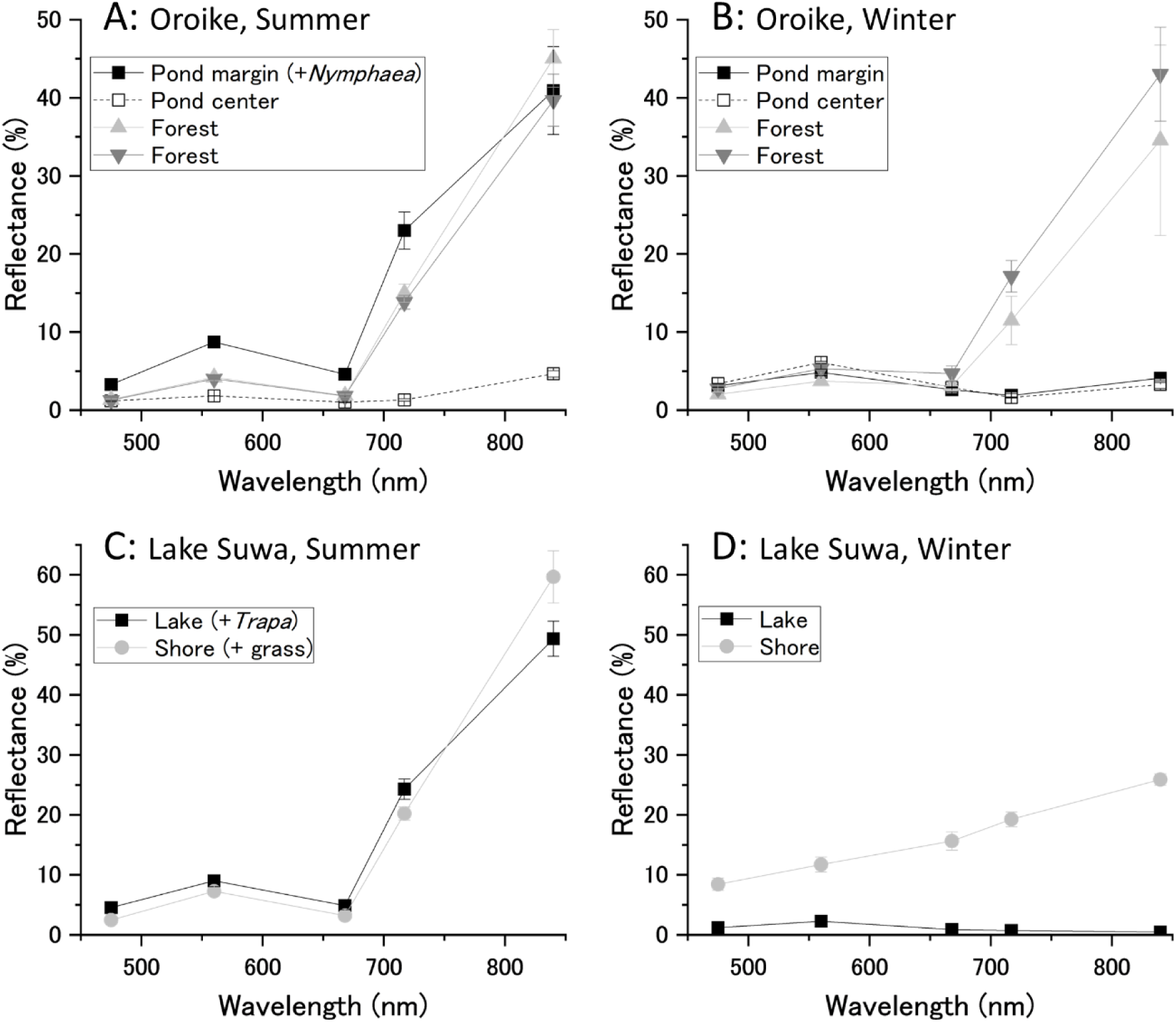
Measurement of reflectance changes exhibited by natural floating vegetation. Spectral reflectance of floating vegetation in a small pond (Oroike) and a relatively large lake (Lake Suwa) was measured by aerial photography using a drone. **A.** The reflectance of vegetation in the pond (Oroike) and its surrounding forest in summer was compared. Water lilies (*Nymphaea spp.*) were predominantly located along the pond’s edge. The graphs show the reflectance of a densely leaved area (▪), a leafless central area (□), and two forest areas around the pond (▴, ▾). The mean reflectance and standard deviation for the six plots in each area are shown. **B.** The reflectance in the same study site and pond as in A was measured in winter disappearing floating leaves of the water lily population. **C.** The reflectance of flourishing vegetation (*Trapa japonica*) in the lake (Lake Suwa) was measured in summer. The graphs represent the reflectance of a dense floating vegetation area (▪, n = 30 ± sd) and a grassland area on the lake shore (●, n = 10 ± sd). **D.** The reflectance in the same study site and lake as in C was measured in winter. Floating vegetation had entirely disappeared, and the water surface was covered with thin ice.

Figures 4C and 4D present the reflectance spectra of vegetation in a larger water body. In summer (Figure 4C, S7C), large colonies of floating plants, dominated by *Trapa japonica*, formed extensive areas a few kilometers long. These floating plants exhibited significant red-edge features, although slightly less pronounced than those of the surrounding grassland, which was dominated by *Phragmites* and Carex species. During the winter months (Figure 4D, S7D), the floating plant community disappeared completely. At the time of measurement, the lake surface was covered with a thin layer of ice (a few centimeters thick). Although it is technically challenging to synthesize an orthomosaic image of an empty water surface, we succeeded in making a reflectance map of the winter lake because of the thin ice layer with small cracks and irregularities. Water temperatures often dropped to near 0°C, killing most of the submerged plants and algae. Consequently, NIR reflectance decreased to levels lower than red reflectance as water and ice exhibit greater absorbance in the NIR compared to the visible light. Although the terrestrial grasslands also died off in winter, the NIR reflectance remained relatively high due to fallen leaves and leaf litter in addition to soil reflectance.

Comparing the sensitivity of various vegetation indices to water surface coverage by *Trapa japonica* confirmed the same trends observed in the water tray trials: DVI increased linearly with increasing coverage and was high when leaf overlap was significant, while RVI and NDVI were more sensitive to low plant coverage. NDVI values exceeded 0.6 even when only a few percent of the water surface was covered by floating plants (Figure S8).

### 3.4 Satellite remote sensing of freshwater ecosystem

In the previous sections, we measured the light reflectance of individual leaves, populations, and portions of natural vegetation. The characteristics of floating vegetation suggest that high-sensitivity detection can be achieved using NDVI as an indicator, based on its contrast with the water surface reflectance. In particular, in lakes of a certain size and depth, where vegetation is eliminated due to freezing temperatures, NIR reflectance becomes lower than red reflectance (Figure 4D), leading to negative NDVI values. Even with water surface coverage below a few percent, we can observe cyclic variations with high sensitivity, as NDVI values switch between positive and negative during the summer and winter months. In the following sections, we assess whether the growth and decline of floating vegetation can be detected from reflectance measurements of entire lakes. To achieve this, we analyzed the NDVI time series data of 72 lakes using clustering techniques. This classification facilitated a detailed examination of NDVI seasonality attributable to floating vegetation. Specifically, 23 locations were classified into Cluster 1 (Figure S9A), 35 locations into Cluster 2 (Figure S9B), and 14 locations into Cluster 3 (Figure S9C).

Lake Suwa (Figure 5, dotted line, and Lake ID 114 in Figure S9B), where we observed *Trapa japonica* colonies, was classified in Cluster 2. Mikata Lake (Lake ID 108), also known to be predominantly covered by *Trapa* populations during summer (Kato et al., 2016), was similarly classified in Cluster 2. The reconstructed seasonality of lakes in this cluster exhibited minimum NDVI values around DOY 90 or 340, corresponding to late February to early March or December, and peak values around DOY 250, which falls in early to mid-September. This seasonal variation clearly reflects the vegetation decline during winter and peak growth in summer. Some lakes showed notable fluctuations in NDVI values, with amplitudes exceeding 0.6 over the course of the year (e.g., Lake ID 69 in Figure S9B). High summer NDVI values in these shallow marshes are likely influenced more by emergent plants, such as lotus, than by free-floating plants. In the present analysis, which examines the detectability of floating plants as a contrast to terrestrial plants, we categorize all aquatic plants that develop their leaves above the water surface as “floating plants” as already explained. These large amplitude fluctuations suggest significant dynamics within aquatic ecosystems, distinguishing them from terrestrial ecosystems.

**FIG. 5.**
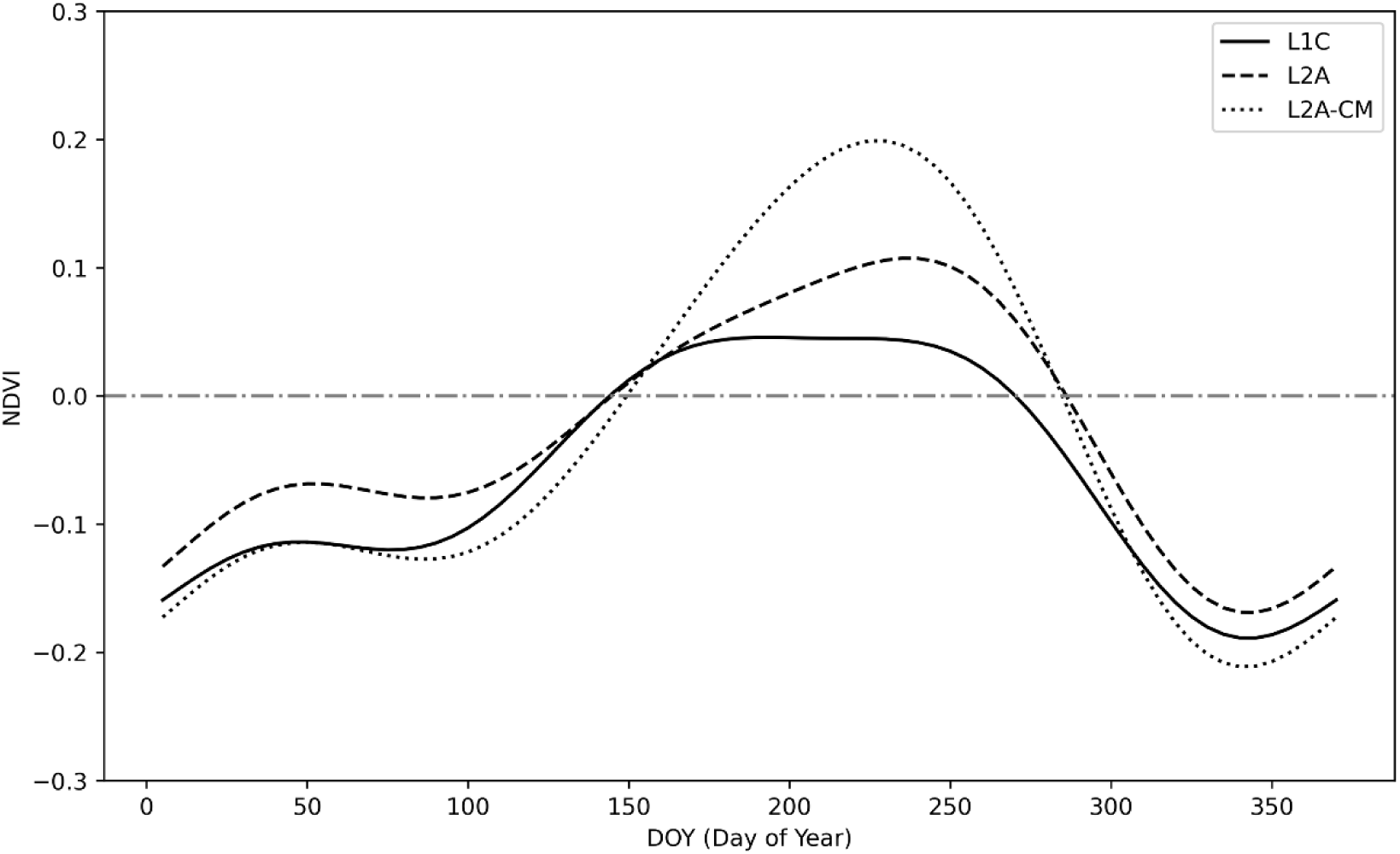
NDVI time series variation based on DOY for Lake Suwa (Lake ID 114 in Figure S7B). The solid line (━) represents the L1C product, the dashed line (---) represents the L2A product, and the dotted line (‥‥) represents the L2A-CM product.

### 3.5 Effect of cloud cover on the NDVI detection

The NDVI analysis in the previous section excluded cloud-covered areas from the satellite images and reflected measurements taken under clear skies. In exoplanet observations, however, the global reflection is measured as a single pixel, making it impossible to exclude cloud effects in the same manner. To provide a more realistic evaluation of vegetation detection for exoplanet observations, we assessed the impact of cloud cover and atmospheric conditions on NDVI detection.

Stratified data were analyzed for lakes belonging to Cluster 2, identified as having typical floating vegetation in the previous clustering analysis. Results using Lake Suwa as an example are shown in Figure 5. The maximum NDVI (NDVI_Max_) values obtained from L2A-CM (Figure 5, dotted line), L2A (Figure 5, dashed line), and L1C (Figure 5, solid line) were 0.20, 0.11, and 0.05, respectively. This indicates that NDVI_Max_ was attenuated by 45% (with 55% remaining) due to cloud cover, and further decreased by 54% (with 46% remaining) due to atmospheric molecules, resulting in a total attenuation of 75% (with 25% remaining). When comparing the magnitude of the difference between maximum and minimum NDVI values (ΔNDVI), the values for L2A-CM, L2A, and L1C were 0.41, 0.28, and 0.23, respectively. The total signal attenuation effect was calculated at 44% (with 56% remaining). Cloud cover slightly shifted the peak position of the seasonal variation curve, while additional atmospheric effects further flattened the peak. Despite these changes, the periodic pattern of NDVI variation between negative and positive values in winter and summer was maintained.

We extended this analysis to all sites in Cluster 2 to confirm whether these signal attenuation effects are common in the detection of aquatic vegetation. The median values of NDVI_Max_ for L2A-CM, L2A, and L1C were 0.205, 0.116, and 0.075, respectively (Figure 6A). The effect of clouds and atmospheric gasses were estimated at 43.4% and 35.3%, respectively. The median values of ΔNDVI were 0.302, 0.167, and 0.129 for L2A-CM, L2A, and L1C, respectively, with cloud and atmospheric gas effects estimated at 44.7% and 22.8%. The total attenuation effect was 63.4% for NDVI_Max_ and 57.2% for ΔNDVI. Although the variation among locations is too large to draw definitive statistical conclusions, ΔNDVI appears to be slightly less susceptible to attenuation effects than NDVI_Max_.

**FIG. 6.**
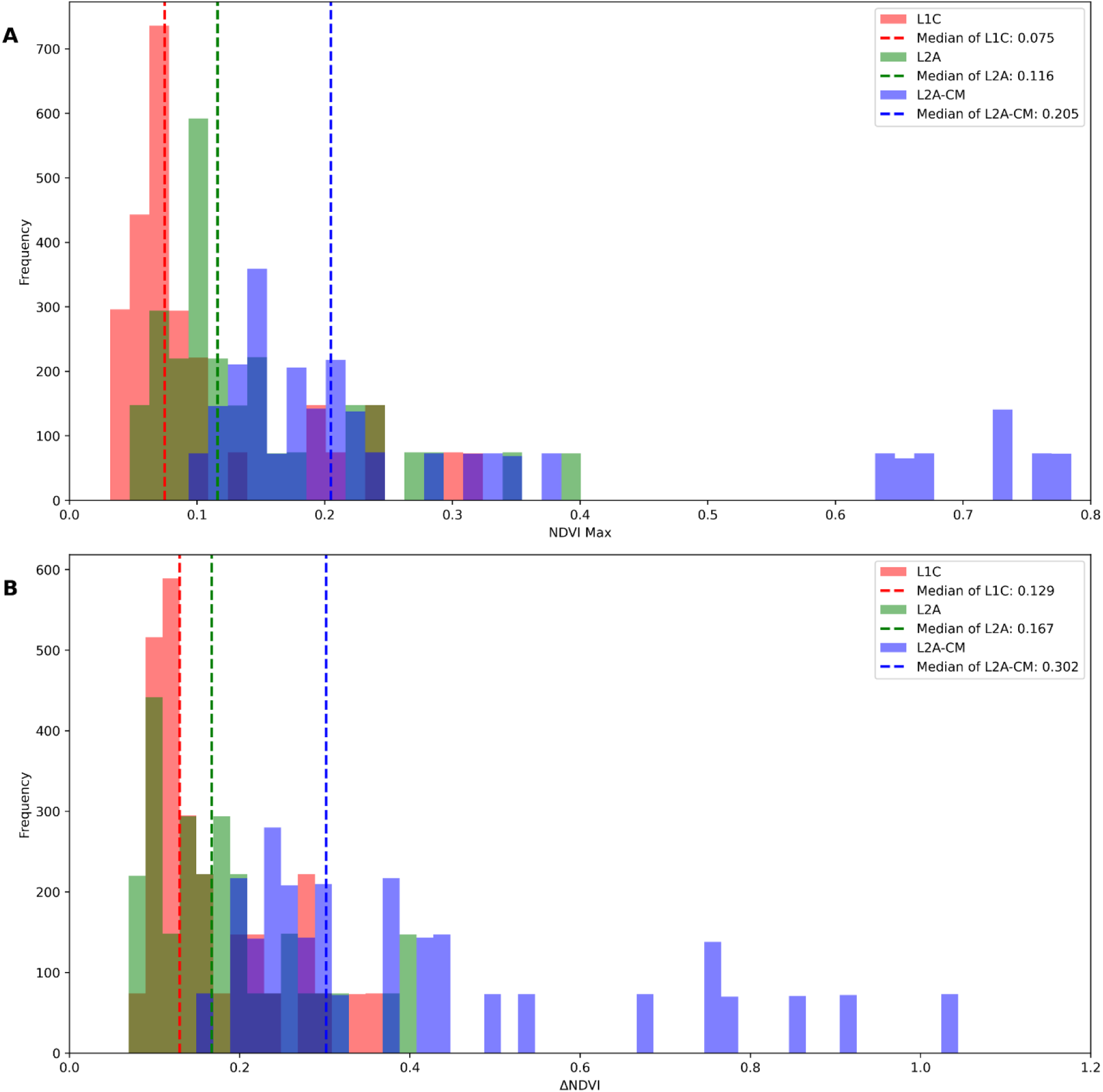
Signal attenuation effects of clouds and atmospheric molecules on NDVI-related indicators. The maximum NDVI value (NDVI_Max_) and the range of variation (ΔNDVI) were calculated from the HANTS-fitted data in Cluster 2, as shown in Panels **A** and **B**, respectively. The red histograms represent the analysis results for the L1C dataset, which includes cloud and atmospheric gaseous effects. The green histograms display the results for the L2A dataset, which includes effects from atmospheric gas only. The blue histograms show the results for the L2A-CM dataset, which excludes cloud and atmospheric gaseous effects. Dashed lines indicate the median values for each histogram.

## 4. Discussion

### 4.1 Spectral reflectance properties of floating plants

Our laboratory-scale reflectance measurements confirmed a pronounced VRE in floating plants, which was not evident in submerged aquatic plants and algae. The green alga *Chlamydomonas reinhardtii*, which is evolutionarily related to terrestrial plants and belongs to the same green lineage, shares the same pigment composition and photosynthetic machinery as vascular plants (Erickson et al., 2015; Grossman et al., 2004). Although dense algal water exhibits a reflectance spectrum similar to that of terrestrial plants, its NIR reflectance amplitude is significantly lower due to the absence of leaf tissue structures. Furthermore, the red-edge wavelengths of algal water were blue-shifted by several tens of nanometers, depending on cell density. The red-edge wavelength of algal cells in a pellet state is approximately 700 nm (data not shown), which is not significantly different from that of terrestrial plants. However, in cell suspensions, the red-edge shifts to shorter wavelengths because reflectance at longer wavelengths is more significantly reduced by water absorption. To accurately predict the reflectance spectra of plants in aquatic environments for future exoplanet exploration, it is essential to consider the attenuation effects due to water, in addition to the inherent reflectance spectra of the plants.

The leaves of floating plants exhibited red-edges at similar wavelengths to those of terrestrial vascular plant leaves (≈710 nm) and similarly high NIR reflectance. The leaf structures required for floating on water may contribute to increased reflectance, as discussed in the following section. However, the high NIR reflectance of individual leaves is partially reduced due to wetting when in contact with the water surface. For floating plants with leaves that rest horizontally on the water surface, the underside of the leaves is inevitably in contact with the water. If the lower surface of the leaf is water-repellent and covered with an air layer, similar to the upper surface, the leaf cannot maintain a stable vertical orientation above the water. Floating plants with horizontal leaves position them above the water surface, without overlap, to optimize light absorption and gas exchange. Therefore, their VRE is smaller than that of terrestrial vegetation with a well-developed canopy structure.

Floating plants with leaves that protrude entirely into the air, such as *Eichhornia crassipes*, eliminate the effects of water and maintain a strong red-edge signal similar to that of terrestrial plants. Additionally, as multiple leaves can be vertically separated, the VRE in these plants is larger than that of floating plants with horizontal leaves. However, unlike terrestrial plants, which can grow several meters tall by anchoring themselves with roots in the ground, floating plants cannot grow more than several tens of centimeters above water, resulting in a smaller VRE.

While contact with water can make VRE detection challenging, the low reflectance of water from visible to NIR wavelengths provides a low-noise background, facilitating the remote detection of VRE. Even when vegetation density is low and water surface exposure is high, the observed VRE largely reflects the signal from the vegetation. This effect is particularly evident when using NDVI as an indicator of VRE. An empty water surface produces negative NDVI values, which are converted to positive values in the presence of even a small amount of floating vegetation. Although NDVI may not correlate linearly with the biomass of vegetation on water, it is highly sensitive to the presence of vegetation, presenting both challenges and opportunities for detection.

Our trials with water trays and observations of natural floating vegetation from low altitudes demonstrated that NDVI can detect vegetation with high sensitivity. However, the absolute value of NDVI was not always a reliable indicator in larger-scale trials, where the entire lake was treated as a single pixel in satellite image analysis. Some small marshes had peak NDVI values of up to 0.5, while relatively large ponds had maximum NDVI values of about 0.2 or less, which is lower than the typical forest value of 0.8 (Huang et al., 2021; Lastovicka et al., 2020). The distribution of floating leaves from emergent aquatic plants is usually confined to areas near the lakeshore at depths of less than a few meters (Lacoul and Freedman, 2006). Free-floating plants can occupy broader areas but are often found in high density along the lakeshore, influenced by wind and water currents.

Therefore, we aimed to detect vegetation-derived periodicity in satellite images by averaging very low-resolution reflectance over the entire lake across an extended period, rather than focusing on high-resolution reflectance during peak growth seasons. We conducted a comprehensive analysis of 148 lakes and marshes in Japan and successfully detected the seasonal NDVI cycles attributable to floating vegetation. Although cloud cover did affect vegetation detection by satellite remote sensing, the impact was manageable, with signal intensity attenuating roughly in proportion to the level of cloud cover.

Specifically, both NDVI_Max_ and ΔNDVI experienced about 45% attenuation due to cloud effects, which corresponds approximately to cloud coverage. Despite this attenuation, the distinct seasonal variation associated with floating vegetation remained evident, demonstrating the robustness of this detection method. When additional atmospheric effects were considered, the attenuation of NDVI_Max_ and ΔNDVI increased to 63% and 57%, respectively, causing greater distortion of the seasonal pattern. ΔNDVI, which measures the difference between two points in the seasonal cycle, was less affected by attenuation than NDVI_Max_, which is based on a single point. This suggests that analyzing the overall pattern of NDVI variation provides a more robust approach for vegetation detection. This methodology could potentially offer a more reliable technique for detecting vegetation in environments with variable atmospheric conditions.

The survey area falls within the temperate (Cfa) and cold (Dfa, Dfb) zones according to the Köppen climate classification (Peel et al., 2007). In tropical regions, where floating vegetation could be denser, seasonal changes may be less pronounced, making maximum NDVI a better indicator. If global topographic and vegetation data comparable to that of the present survey become available, it will enable a more comprehensive evaluation of the observability of floating vegetation.

### 4.2 Morphology and evolution of floating plants

Floating plants have evolved to remain buoyant on the water surface and perform photosynthesis efficiently. They possess surface and internal structures that enhance surface tension (preventing leaves from submerging) and buoyancy (keeping leaves afloat). These features also enhance light reflectance, making the remote detection of floating vegetation easier.

All plants (Embryophytes) produce waterproof compounds and form a cuticle layer on the leaf surface (epidermis) to protect against environmental stress (Post-Beittenmiller, 1996). Cuticular waxes play a crucial role in preventing water loss from the leaf surface to the air. In contrast, plants in humid regions tend to have waxy, superhydrophobic leaves, known as the lotus effect, to prevent water from entering the stomata (Ensikat et al., 2011). Floating plants have waxy leaves to keep their surfaces dry and enhance surface tension in water. The developed cuticle layer increases specular reflection. Additionally, hairs on the leaf surface (trichomes) serve as a physical barrier, protecting leaves from adverse environments (Wang et al., 2021). These microscopic hairs repel water, preventing the leaf from sinking. For instance, the dense hairs found on aquatic ferns (e.g., *Azolla filiculoides, Salvinia molesta*) can form an air layer on the upper leaf surface, even on turbulent water. These dense hairs increase light scattering at the leaf surface, thus enhancing light reflection (see Figures 1Cd and 1Df). Some angiosperm floating plants (e.g., *Nymphaea spp*. and *Limnobium laevigatum*) have thick, spongy tissues with air chambers (Kaul, 1976) to increase buoyancy. Other species (e.g., *Eichhornia crassipes* and *Trapa japonica*) have bulb-like, spongy nodules in their stalks specialized for floating. Light scattering from these internal structures further increases reflection.

These traits of floating plants were acquired after the terrestrial evolution of plants (Gensel, 2008). The invasion of plants from the ocean to land via freshwater began in the early Paleozoic, coinciding with the continental realignment from Gondwana to Pangaea (Dahl and Arens, 2020). The colonization of early terrestrial plants (around 500 million years ago) was initially restricted to wet lowland environments. While some mosses (bryophytes) are riparian or submerged, there is no fossil record showing floating morphologies. After the emergence of vascular plants (approximately 420 million years ago), terrestrial vegetation gradually expanded its habitat to drier uplands. The floating plants found in modern freshwater ponds are classified as ferns and angiosperms, which were the dominant terrestrial plants in both past and present ecosystems. Compared to the conquest of desiccated soil (around 360 million years ago), the readaptation of vascular plants to freshwater environments occurred much later. Fossil records and phylogenomic analyses indicate that aquatic ferns emerged around 145 million years ago (Martín-Closas, 2003; Shen et al., 2018). Aquatic angiosperms are thought to have emerged at the beginning of angiosperm evolution, approximately 140 million years ago (Du et al., 2016; Soltis et al., 2008). The evolutionary sequence of vascular plants suggests that the acquisition of desiccation tolerance facilitated their adaptation to water environments. Once vascular plants acquired essential traits (e.g., cuticulated epidermis and lignified conducting tissue), they were well-prepared to expand their habitat above water, managing to draw water up without wetting the leaf surfaces.

While a diverse array of macrophytes, including photosynthetic bacteria and kelp, thrive in seawater, floating plants predominantly inhabit freshwater environments. Aquatic plants (including floating, emergent, and submerged species) account for less than 1% of vascular plant species (Chambers et al., 2008), and among these, only a small fraction (fewer than 100 species) are adapted to saltwater. Most aquatic plants are equally or more susceptible to salinity stress compared to terrestrial plants (Moreira et al., 2023). Submerged marine angiosperms, such as seagrasses, are adapted to coastal areas (Touchette, 2007), but their mechanisms for salt tolerance are insufficient to allow widespread distribution in the ocean, unlike sea algae.

Although floating plants are limited in both species diversity and habitat, they significantly impact the local environment and ecosystems at the water-land interface (Chambers et al., 2008). Furthermore, it has been suggested that floating plants may have influenced global climate in the past. Sub-seafloor excavations in the Arctic Ocean have revealed 50-million-year-old sediments containing large amounts of Azolla fossils (Brinkhuis et al., 2006). This finding supports the Azolla Event hypothesis, which proposes that *Azolla* blooms covering much of the Arctic Ocean persisted for nearly a million years, absorbing substantial amounts of CO_2_ and depositing it into the seafloor, thereby contributing to a rapid decline in global temperatures. Although there is evidence that challenges this hypothesis (Neville et al., 2019), it remains a valid consideration when examining the coevolution of plants and planetary environments. Notably, it deserves consideration as a model for plant evolution on a water-rich exoplanet.

### 4.3 Aquatic VRE observations on exoplanets

On Earth-like exoplanets with continents dispersed within oceans, terrestrial vegetation could evolve similarly to Earth, with VRE appearing at different evolutionary stages (O’Malley-James and Kaltenegger, 2018a). Floating vegetation might thrive more than on Earth today due to slight variations in surface conditions. Ocean and lake salinities have varied significantly throughout Earth’s history, influenced by processes such as continental weathering, which increases salinity, and salt decomposition in supratidal zones, which reduces it (Knauth, 2005). The size distribution of lakes on Earth, in terms of surface area, follows well-defined scaling relationships governed by geological processes, with smaller lakes being far more abundant than larger ones (Cael and Seekel, 2016). These freshwater systems, formed since earlier geological periods, may have played a critical role in shaping early ecosystems and supporting the evolution and diversity of life, forming the foundation for the modern biosphere. The more complex the continental topography and the more active the hydrological cycle, the more likely it is for freshwater caps to form both on land and in the oceans. If an Earth analog supports extensive floating plant vegetation in addition to terrestrial plants, the observed VRE could be more significant than on modern Earth. Assuming that floating plants evolve from terrestrial plants, as observed on Earth, the wavelength position of the VRE in both terrestrial and aquatic vegetation would likely be similar, thus enhancing the detectability of vegetation indices such as NDVI.

O’Malley-James and Kaltenegger (2018a) suggested that hotter, drier planets are more likely to exhibit VRE due to the presence of more plants with high VRE and fewer clouds that could attenuate the VRE signal. However, VRE detection might also be possible on water-rich planets if the environment is conducive to the growth of floating plants. This study shows that NDVI is a highly sensitive indicator for floating vegetation, and its seasonal variation is less affected by cloud cover than its absolute values. The seasonality in NDVI, associated with periodic environmental changes, may be detectable on exoplanets.

Super-Earths and sub-Neptunes in the habitable zone can be covered by oceans several hundred kilometers deep (Nixon and Madhusudhan, 2021). Oceanic planets are among the candidates for life exploration; however, if high water pressure from excess water leads to the formation of ice layers between silicate rocks and surface water, the likelihood of life originating is reduced (Ollivier, 2011). Moreover, it remains unclear whether underwater phototrophs can evolve into floating plants without undergoing terrestrial evolution. If evolution on ocean planets is restricted to macrophytes in water, VRE detection is unlikely. In such scenarios, fluorescence may serve as a more effective biosignature, particularly if abundant biota are present in the water (O’Malley-James and Kaltenegger, 2018b; Komatsu et al., 2023).

## 5. Conclusions

Aquatic plants with floating leaves tend to exhibit a pronounced red-edge among vascular plants, primarily due to their specialized morphology, which is essential for efficient photosynthesis on the water surface. This morphology, associated with the red-edge feature, is likely to be expected even if these plants evolved along different evolutionary paths than those from vascular plants. At the community scale, floating vegetation that spreads horizontally with minimal leaf overlap displays a less pronounced red-edge compared to terrestrial vegetation, which often grows vertically with multiple overlapping leaves, thereby enhancing the red-edge effect. Although water can reduce the overall reflectance of plants, the low background noise due to the low reflectance of water enhances the detection of floating vegetation. Among the three vegetation indices—RVI, DVI, and NDVI—NDVI was identified as the most effective tool for detecting floating plants, especially in areas where vegetation coverage is low or varies significantly. Our satellite remote sensing analysis demonstrates that NDVI can sensitively detect the reflectance and its seasonal variation in lake-wide floating vegetation, with observed NDVI fluctuations of approximately ±0.2 around zero. This advantage in vegetation detection is generally applicable across different environments, assuming similar water composition and periodic environmental fluctuations. Additionally, although cloud cover can cause approximately 45% attenuation in NDVI values (both NDVI_Max_ and ΔNDVI), our analysis of seasonal NDVI cycles remains effective for detecting floating vegetation under varying atmospheric conditions.

Our research has shown that the red-edge, previously considered an indicator specific to terrestrial vegetation, can also be detected in vegetation over water. This finding has significant implications for future exoplanet surveys, suggesting that floating plants could be a viable target in the search for life on water-rich exoplanets. To produce a detectable signature from exoplanets, exo-floating plants would need to acquire morphology that improves surface tension and buoyancy. These fundamental traits would likely be acquired after plants invade land on an Earth-like terrestrial planet. On an oceanic planet, it remains uncertain whether plants can develop such traits while remaining in the water. Although floating plants currently inhabit local freshwater environments on Earth, their global proliferation in oceans could be conceivable on an exoplanet, as illustrated by the Azolla Event hypothesis.

## Acknowledgments

This work was supported by the Model Plant Section in NIBB. We thank Dr. Kaori Kohzuma for providing the hyperspectral camera and for her help with the spectral imaging and analysis. We also thank Dr. Osamu Watanabe for lending his expertise in vegetation remote sensing.

## Authorship confirmation/ contribution

AM is mainly responsible for spectral reflectance measurements of green algae and satellite remote sensing and authored the relevant sections in the manuscript. KT is mainly responsible for the spectral reflectance measurements of aquatic plants and authored the relevant sections in the manuscript. KT and YK contributed to the conception of this project. All authors contributed to revising the manuscript and approved the submitted version.

