## Supplemental Materials for "Remote Detection of Red-Edge Spectral Characteristics in Floating Aquatic Vegetation"

**FIG. S1** Photographs of plants used for reflectance measurements. The left column shows close-up images of leaves to highlight the differences in leaf morphology. The middle and right columns show larger-scale pictures of the whole plant and its population. The close images for *Lemna minor* and *Azolla filiculoides* were acquired using a digital microscope camera (MC120HD/M165FC, Leica), while other photographs were taken with a standard digital camera.

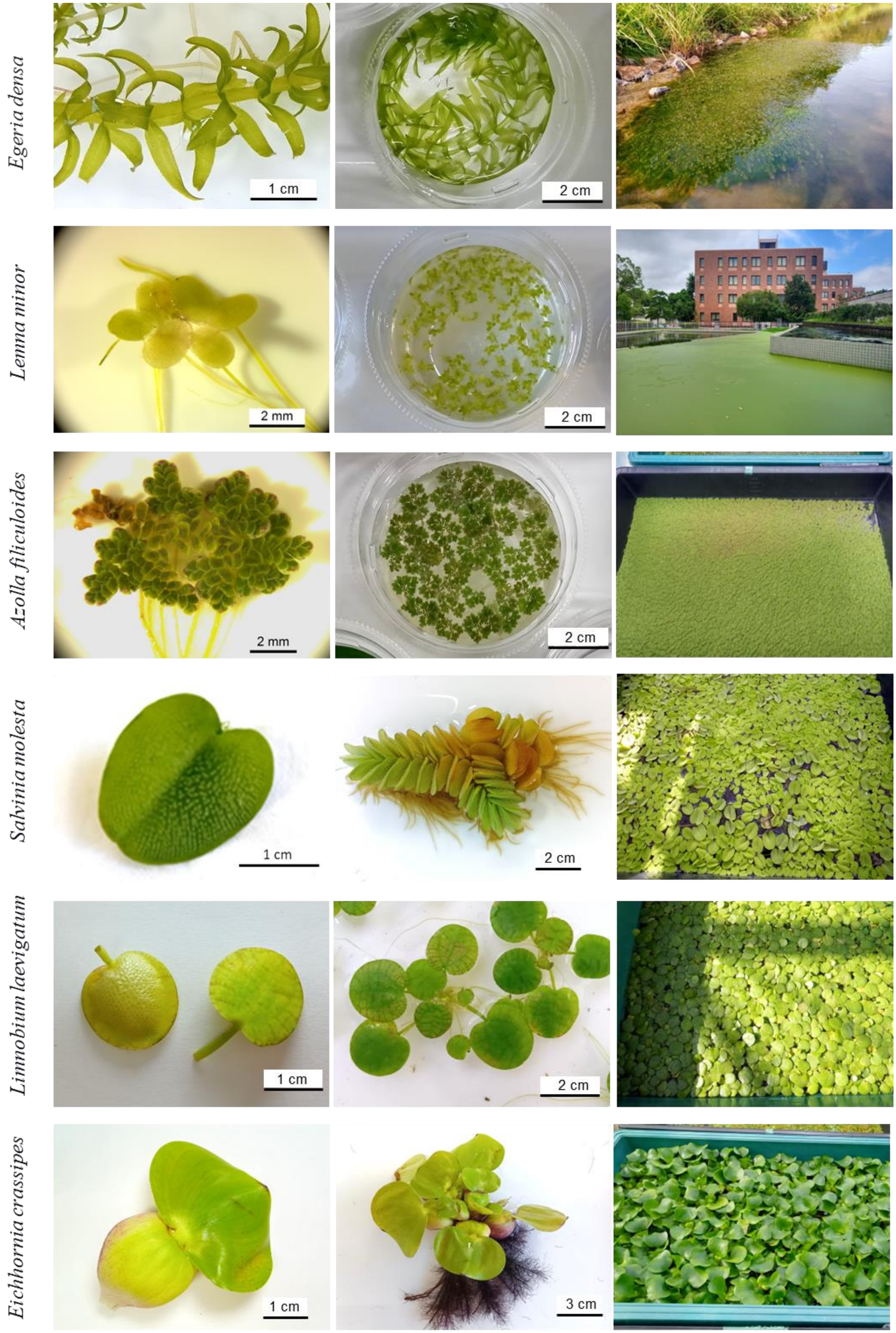
Egeria densa
1 cm
2 cm
Lenna minor
2 mm
2 cm
Azolla filiculoides
2 mm
2 cm
Salvinia molesta
1 cm
2 cm
Limnobium laevigatum
1 cm
2 cm
Eichhornia crassipes
1 cm
3 cm

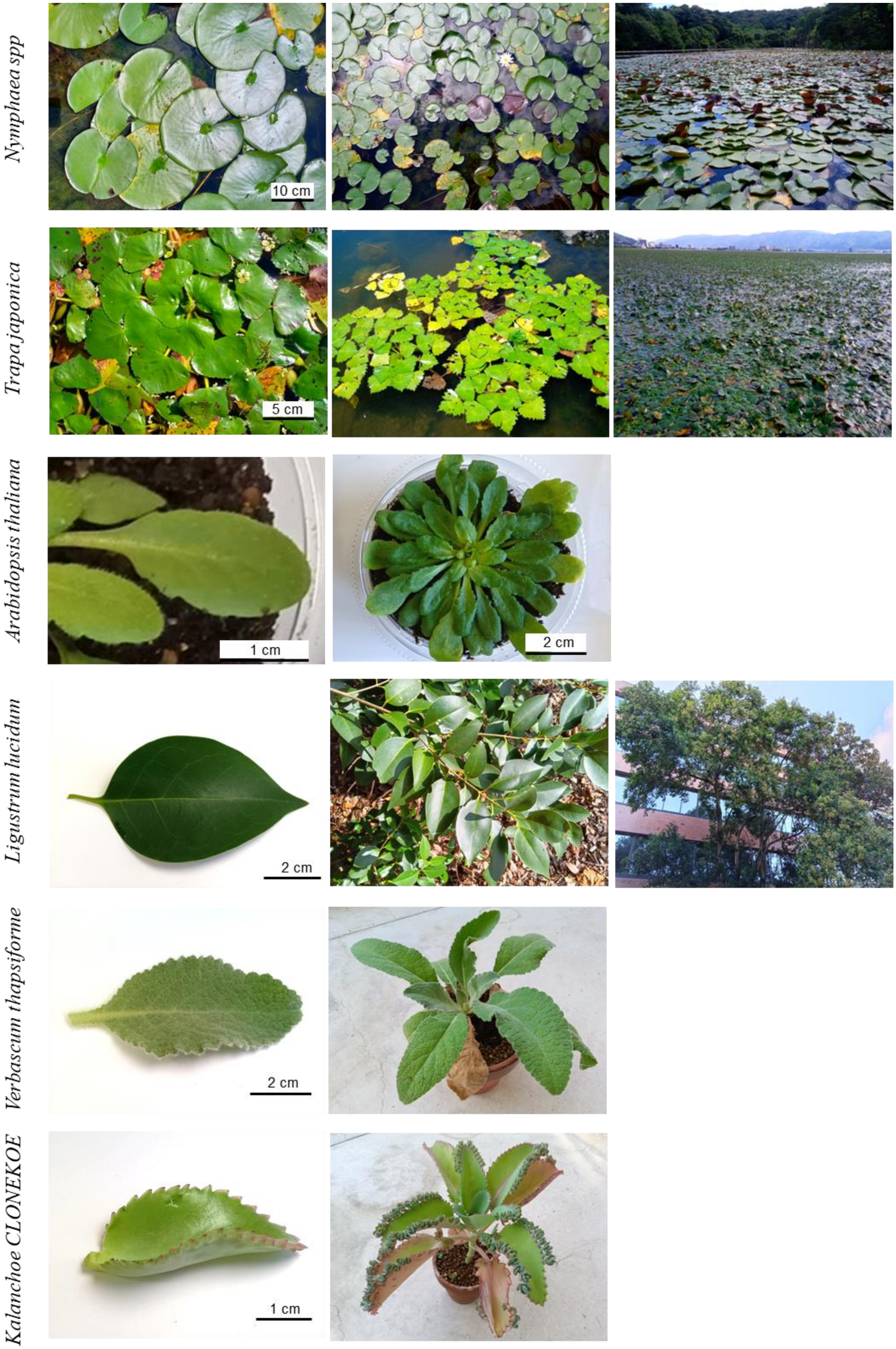
Nymphaea spp
10 cm
Trapa japonica
5 cm
Arabidopsis thaliana
1 cm
2 cm
Ligustrum lucidum
2 cm
Verbascum thapsiforme
2 cm
Kalanchoe CLONEKOE
1 cm

**FIG. S2.** Spectral imaging of aquatic plants using a hyperspectral camera. Spectral reflectance images of aquatic plants were acquired by a hyperspectral camera (KD-1; EBA Japan Co., Ltd.). The camera positioned 30 cm above the sample had a field view of 18 cm x 18 cm (black square in Panel A). Light reflectance was scanned between 400 nm and 900 nm with 5 nm intervals for the central area of 9 cm x 9 cm (160,000 pixels), as shown in Panel A. The reflectance averages were obtained for a central area of 3.6 cm x 3.6 cm (25,600 pixels), where the plants were uniformly distributed (Red square in Panel A and B). RGB image was constructed by reflectance at 680, 550, and 480 nm. Panels A, B, and C are RGB images of *Eeria densa*. Panel D is RGB images of *Azolla filiculoides*. The change in the apparent RGB image when the maximum value of reflectance to be displayed varies from 250 to 4000 is shown in Panels C and D. The interspecies comparison of the RGB images shown in Figure 1a was made using the display settings that resemble the naked eye observation under room light (green boxed images in Panels C and D).

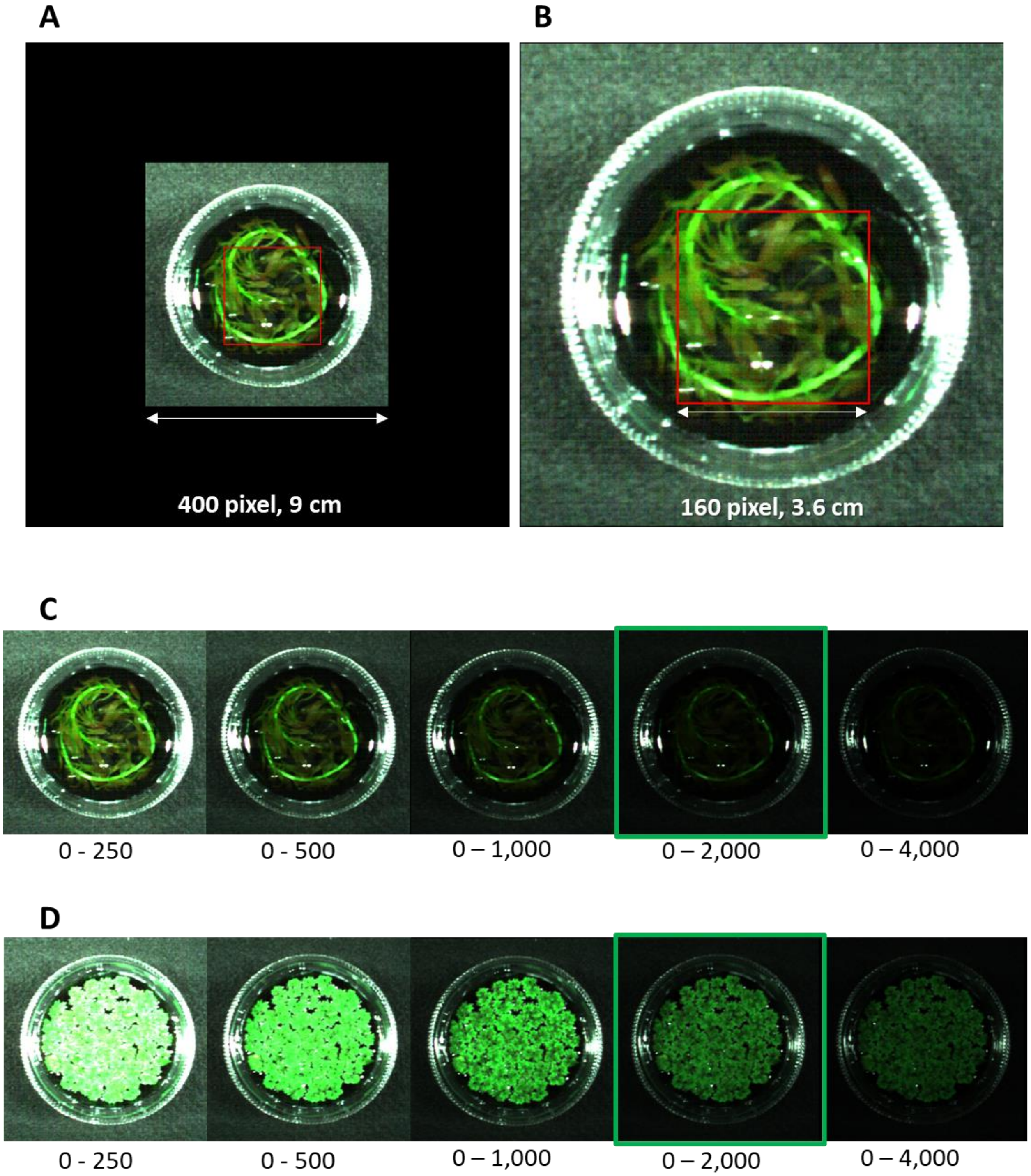
A
B
400 pixel, 9 cm
160 pixel, 3.6 cm
C
0 - 250
0 - 500
0 – 1,000
0 – 2,000
0 – 4,000
D
0 - 250
0 - 500
0 – 1,000
0 – 2,000
0 – 4,000

**FIG. S3.** Spectral imaging of floating plant populations using a multispectral camera. Panel **A** shows five species on water trays (85 x 55cm). Plants were placed at the maximum density to cover the water surface completely. Panel **B** shows *Salvinia molesta* at different densities. Healthy plants were initially packed densely on the surface of the water tray, ensuring no gaps between them. Subsequently, half of them were removed and redistributed evenly across the entire water surface. The relative plant density was adjusted to four levels (1, ½ , ¼, and ⅛). Panel **C** shows 5-band spectral reflectance images for *Salvinia molesta* using RedEdge-M (MicaSense). 475 nm (Blue), 560 nm (Green), 668 nm (Red), 717 nm (RE), and 840 nm (NIR) reflectance of different densities of Salvinia molesta are shown.

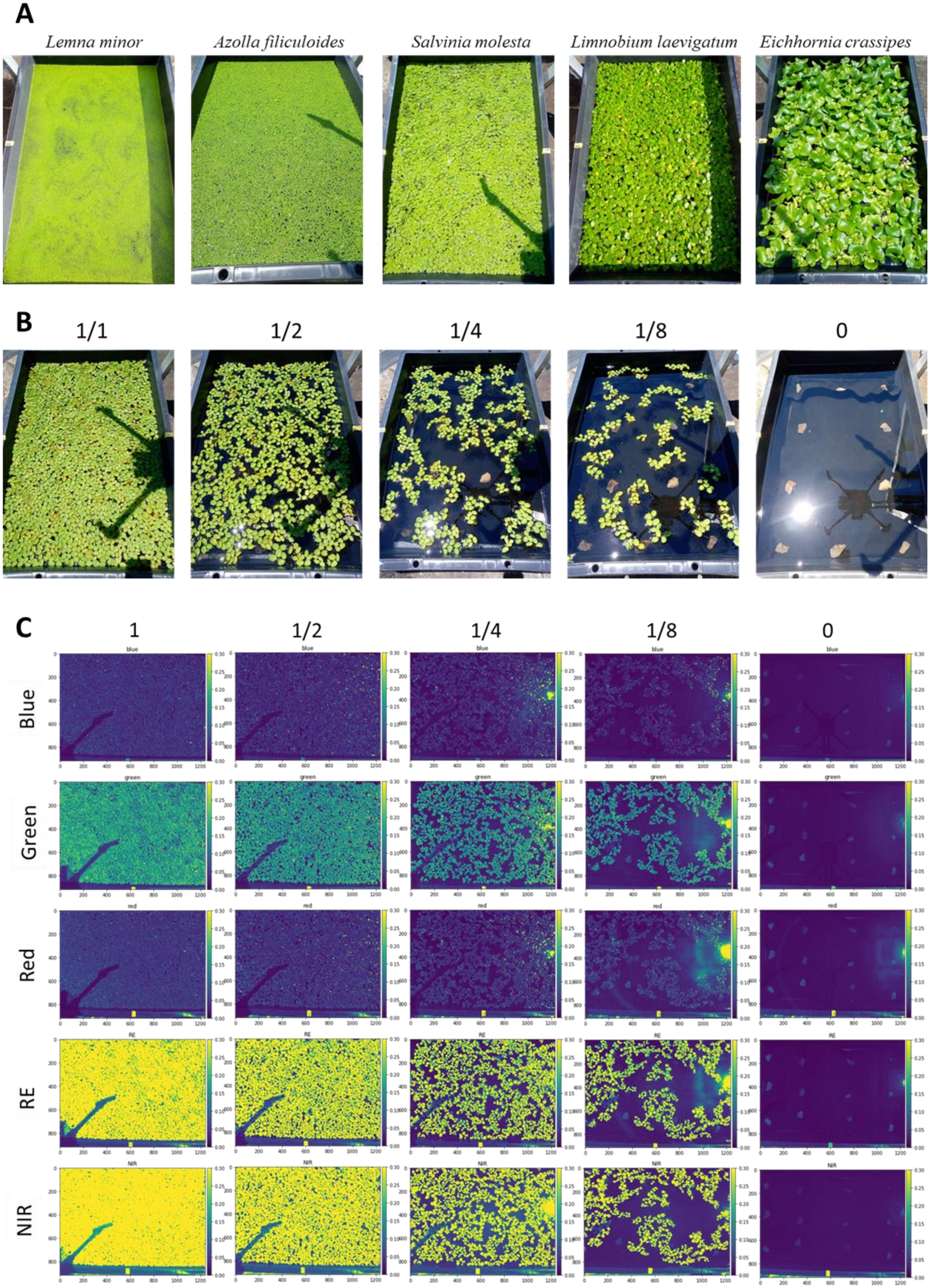
A
Lemna minor
Azolla filiculoides
Salvinia molesta
Limnobium laevigatum
Eichhornia crassipes
B
1/1
1/2
1/4
1/8
0
C
1
1/2
1/4
1/8
0
Blue
Green
Red
RE
NIR

**FIG. S4.** Elbow method analysis for determining the optimal number of clusters in K-shape clustering. The random state parameter was set to 42 for reproducibility. Based on the analysis of the elbow curve, the optimal number of clusters was determined to be three.

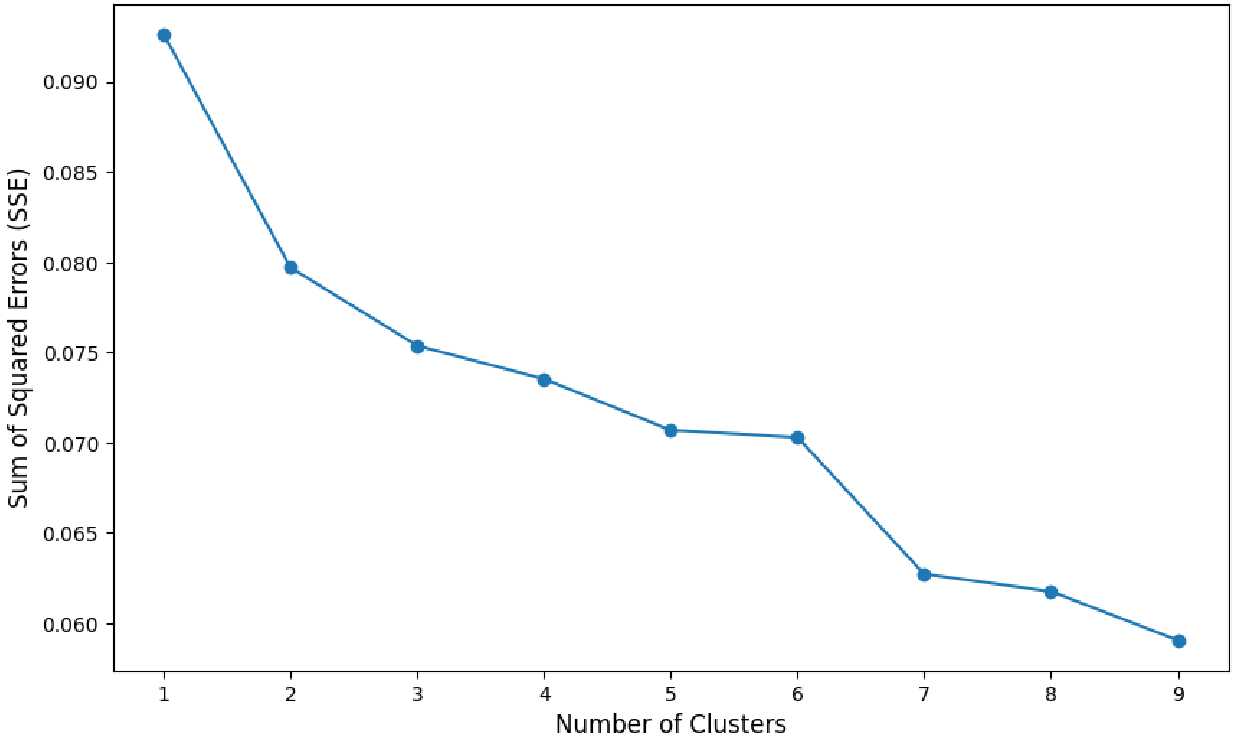

**FIG. S5.** Reflectance spectra of individual leaves. The mean and standard deviation of the five leaves measured are shown by the black line and gray band. The NIR reflectance values of Arabidopsis (35%) are shown as dashed lines in all graphs for comparison.

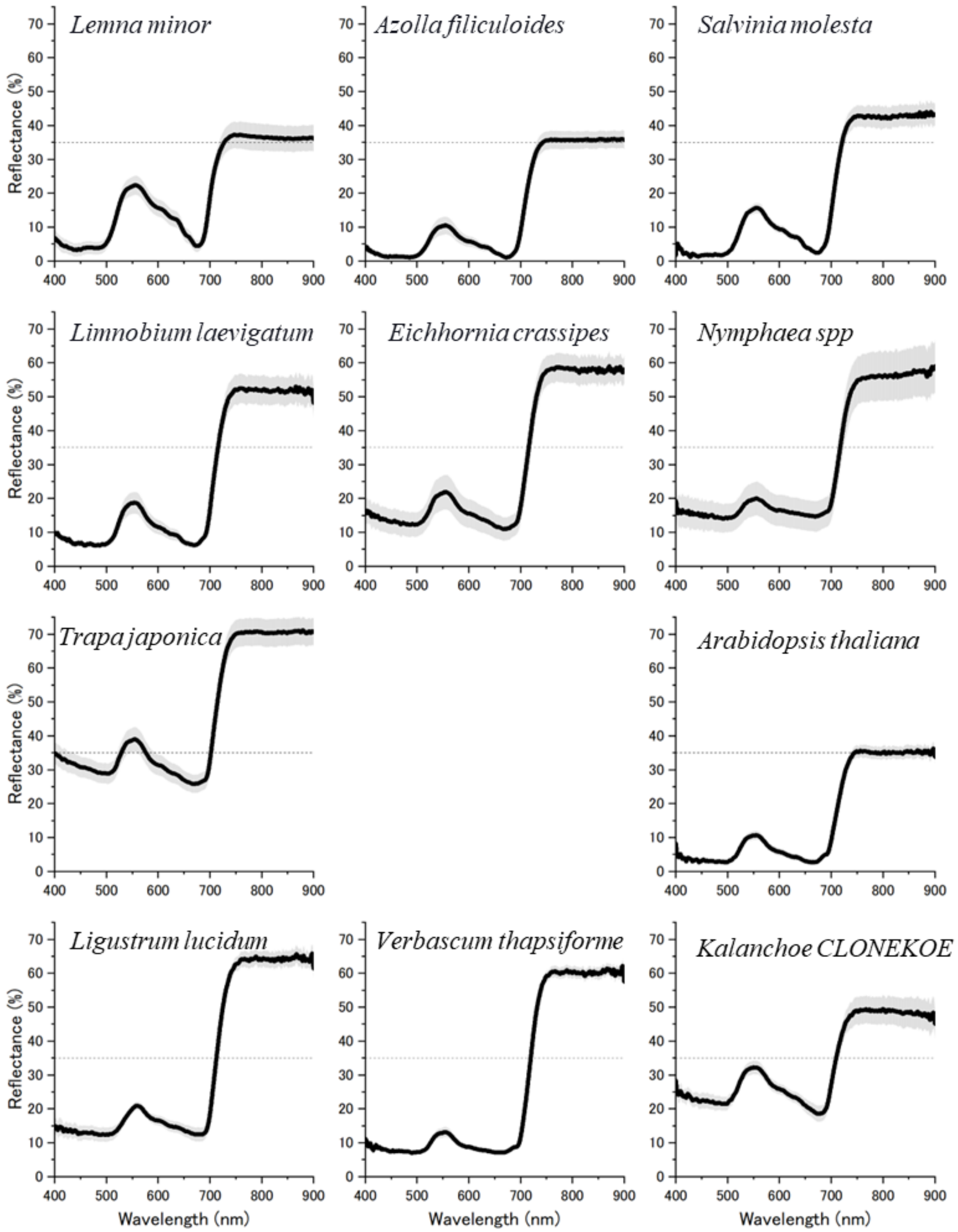

**FIG. S6.** The reflectance spectrum of floating plant populations. Spectral reflectances of 5 species were calculated from 5-band images as described in Figure S3. Each symbol represents the averages from the 3-repeated image acquisitions.

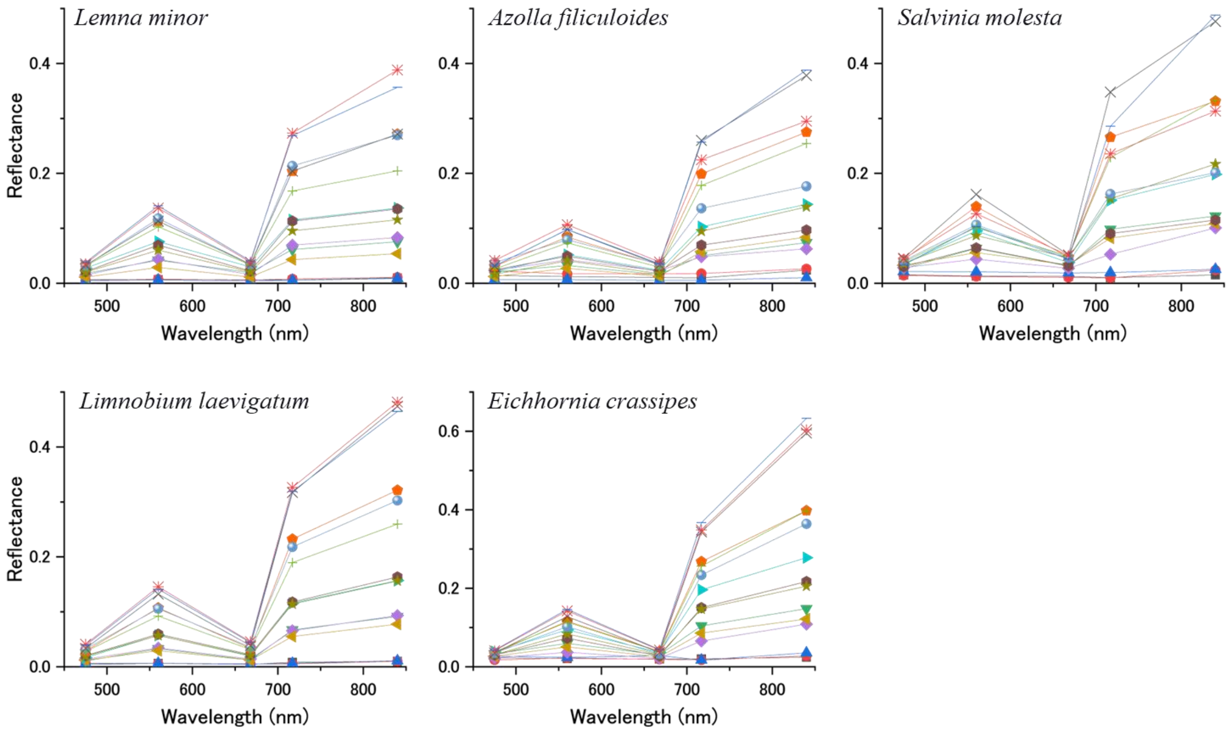

**FIG. S7.** Spectral imaging of natural floating vegetation using a drone-mounted multispectral camera. Five-wavelength orthomosaic images were synthesized from a series of images taken from a drone flying at an altitude of 100 m. Color images were synthesized from red, green, and blue reflectance images. The reflectance of the vegetation and water surface was calculated from the area indicated by the 1-meter squares and is shown in Figure 4. **A** is a color image of water lily (*Nymphaea spp*) vegetation in Oroike in the summer (August 5, 2021). **B** is a color image of Oroike in winter (January 20, 2023). **C** is a color image of water chestnut (*Trapa japonica*) vegetation in Lake Suwa in the summer (September. 13, 2021). **D** is a color image of Lake Suwa in the winter (January 16, 2022).

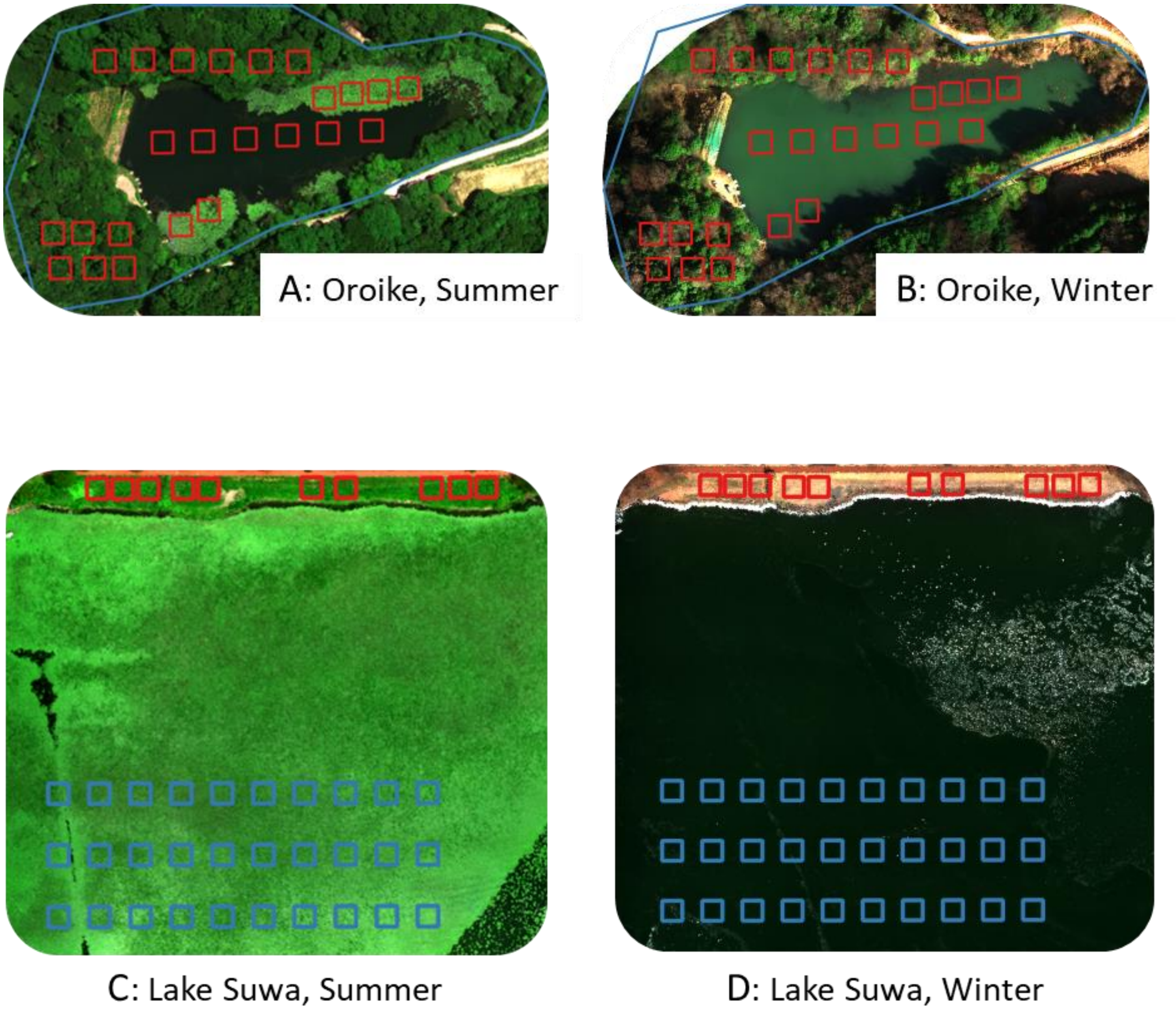

**FIG. S8.** Relationship between vegetation indices and water coverage by *Trapa japonica*. The study site in Lake Suwa was selected at the outer edge of the floating vegetation, where about half of the water surface was covered by floating leaves (July 29, 2022). The 100 x 250m study area was divided into 250 sections with a 10 x 10m mesh. The NDVI grayscale image was binarized with a threshold of 0.2 (A). The cover ratio was calculated for each section and plotted against vegetation indices. B, C, and D show RVI, DVI, and NDVI, respectively.

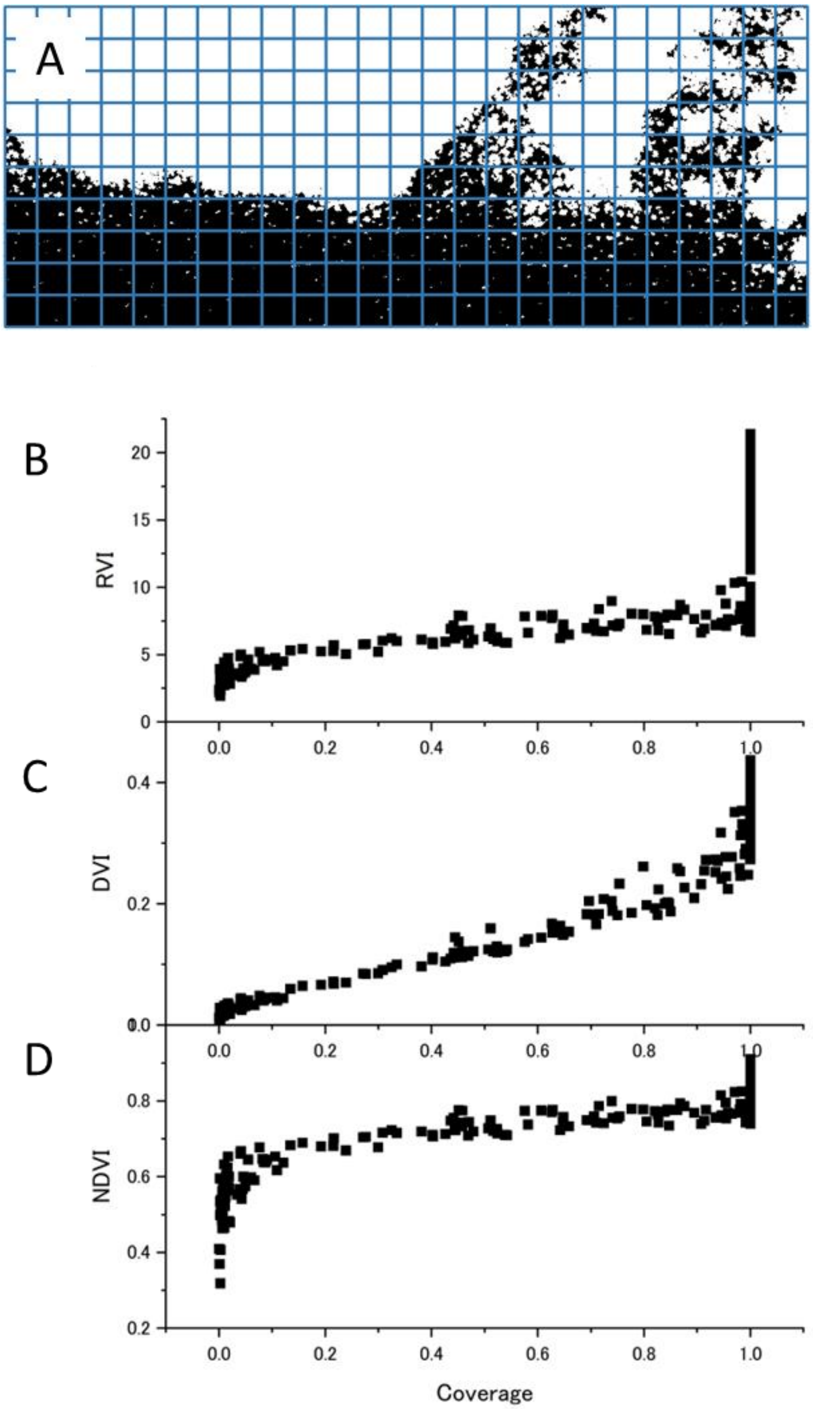

**FIG. S9A.** Annual variation of Cluster 1 from K-shape clustering of NDVI time series data post-HANTS processing. The dashed line indicates the threshold where NDVI equals zero.

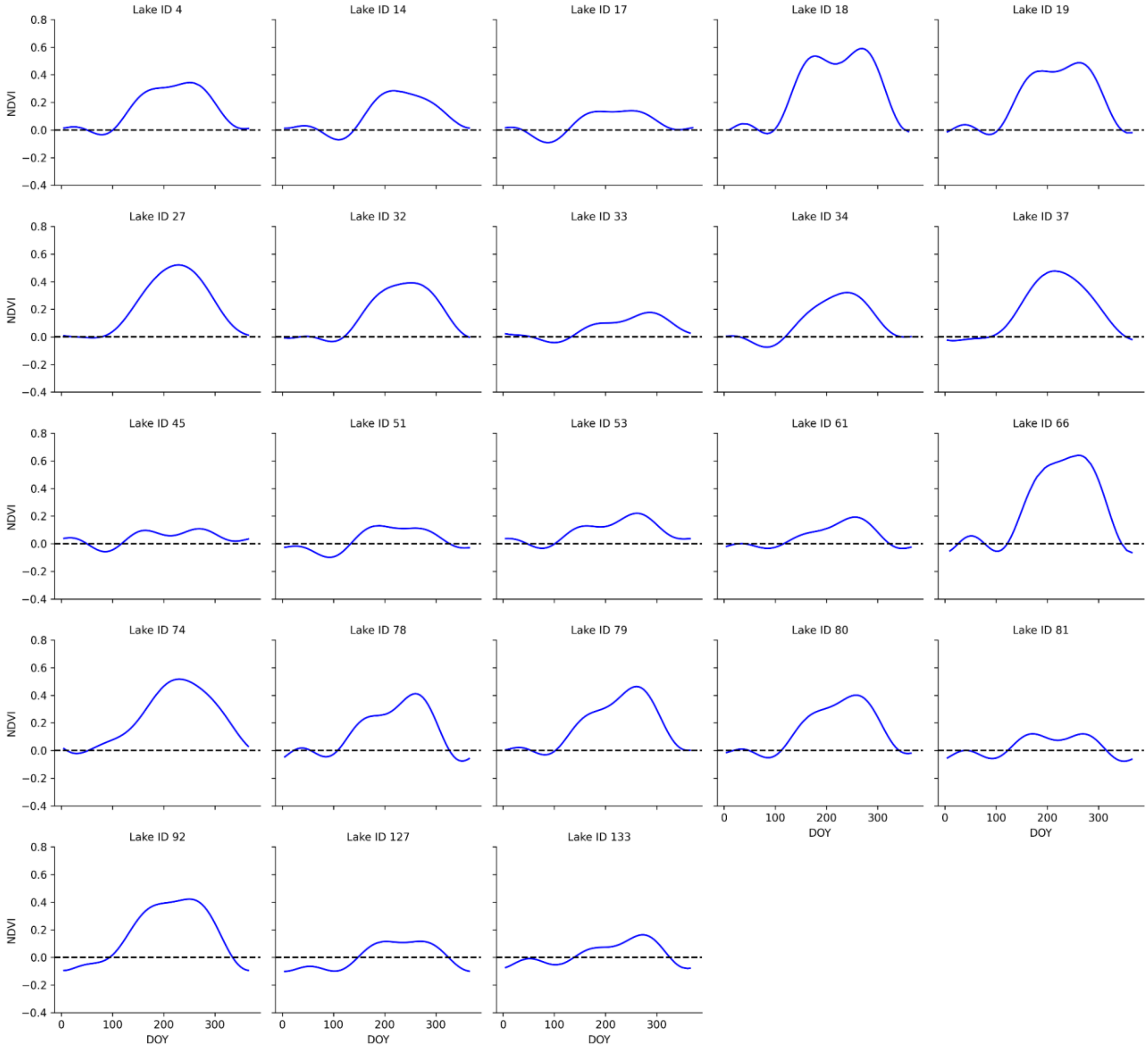

**FIG. S9B.** Annual variation of Cluster 2 from K-shape clustering of NDVI time series data post-HANTS processing. The dashed line indicates the threshold where NDVI equals zero.

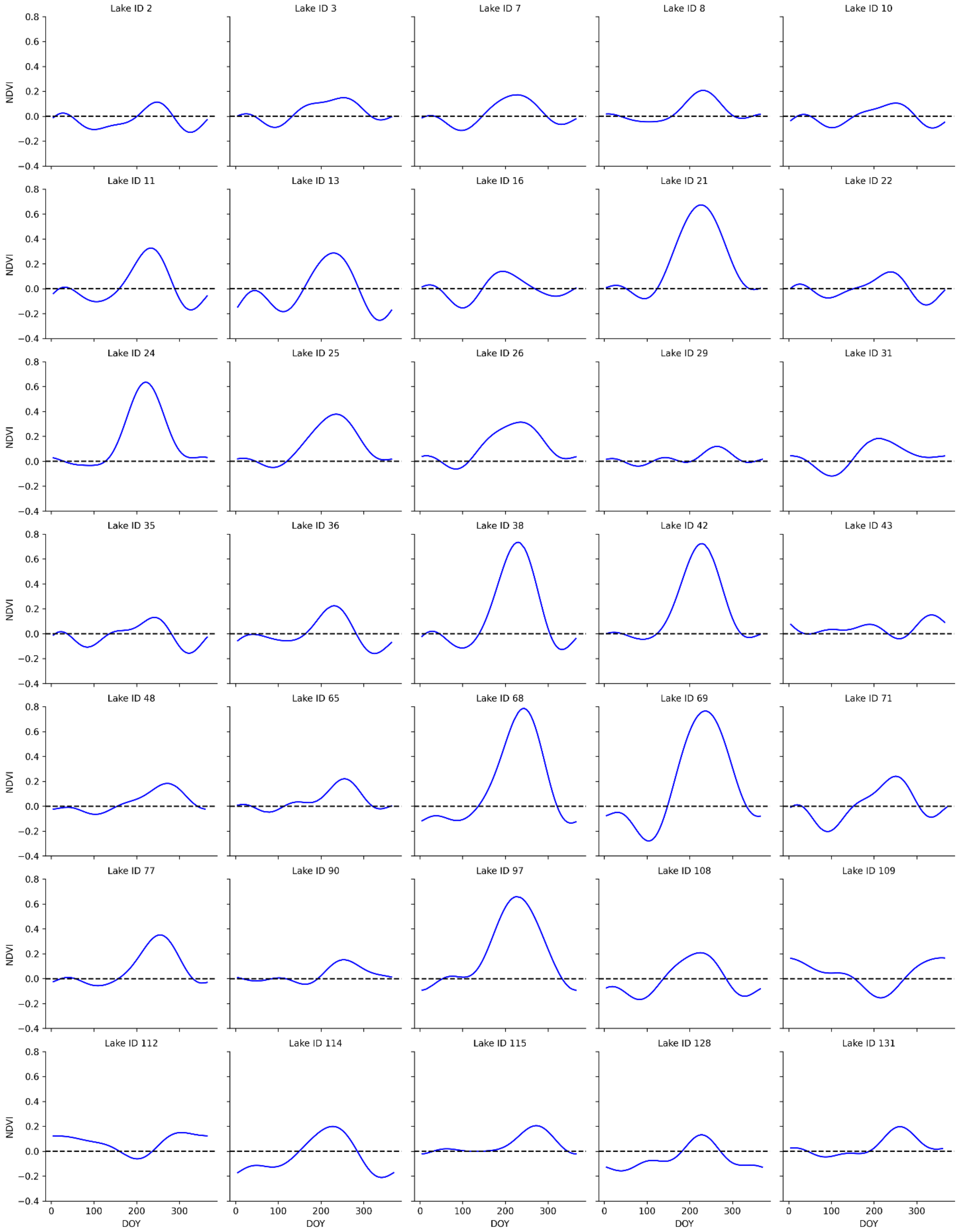

**FIG. S9C.** Annual variation of Cluster 3 from K-shape clustering of NDVI time series data post-HANTS processing. The dashed line indicates the threshold where NDVI equals zero.

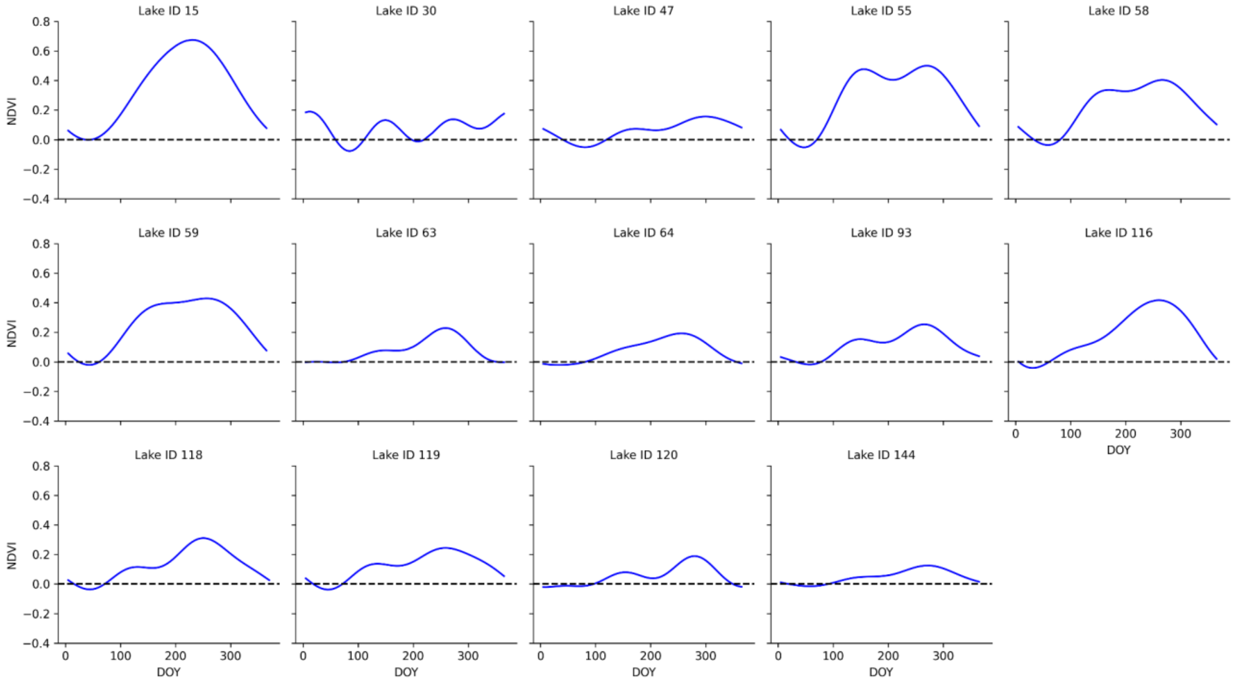

**Table S1.** This table presents the geographic information of the 148 sites analyzed in this study, alongside the NDVI characteristics derived from L2A-CM data and the assigned clustering labels. “Lat” refers to North Latitude, and “Long” refers to East Longitude. "Periodicity" indicates the length of the NDVI cycle in months, with "N/A" representing sites with no clear periodic pattern. "Trend" categorizes the NDVI behavior over the annual cycle: consistently positive values are denoted as (+), consistently negative values as (-), and fluctuating values that alternate between positive and negative as (+/-). "Max" indicates the maximum NDVI value observed throughout the year. Note that the LakeIDs have been explicitly reassigned for this research and do not correspond to the identifiers in the original database.

| | | | | | NDVI | | | |
|---|---|---|---|---|---|---|---|---|
| **Lake ID** | **Lake Name** | **Lat** | **Long** | **Area ($km^2$)** | **Periodicity (Month)** | **Trend** | **Max** | **Cluster No.** |
| **1** | **Harutori** | **42.97** | **144.40** | **0.34** | **12** | **+/-** | **0.02** | **N/A** |
| **2** | **Toufutsu** | **43.93** | **144.40** | **8.19** | **12** | **+/-** | **0.11** | **2** |
| **3** | **Mokoto** | **43.96** | **144.32** | **1.02** | **12** | **+/-** | **0.15** | **2** |
| **4** | **Riyaushi** | **44.00** | **144.16** | **0.42** | **12** | **+/-** | **0.34** | **1** |
| **5** | **Abasiri** | **43.97** | **144.17** | **32.28** | **N/A** | **+/-** | **0.04** | **N/A** |
| **6** | **Notoro** | **44.06** | **144.15** | **58.14** | **12** | **+/-** | **0.05** | **N/A** |
| **7** | **Utonainuma** | **42.70** | **141.71** | **1.93** | **12** | **+/-** | **0.17** | **2** |
| **8** | **Koetoioonuma** | **45.38** | **141.76** | **4.87** | **12** | **+/-** | **0.21** | **2** |
| **9** | **Shibunotunai** | **44.25** | **143.55** | **2.62** | **6** | **+/-** | **0.03** | **N/A** |
| **10** | **Komukenuma** | **44.26** | **143.51** | **3.90** | **12** | **+/-** | **0.11** | **2** |
| **11** | **Chouboshi** | **43.25** | **145.55** | **0.45** | **12** | **+/-** | **0.33** | **2** |

| | | | | | | | | |
|---|---|---|---|---|---|---|---|---|
| **12** | **Fuuren** | **43.31** | **145.33** | **59.10** | **12** | **+/-** | **0.09** | **N/A** |
| **13** | **Onneto** | **43.25** | **145.51** | **5.72** | **12** | **+/-** | **0.29** | **2** |
| **14** | **Okotanpenuma** | **42.80** | **141.26** | **0.41** | **12** | **+/-** | **0.28** | **1** |
| **15** | **Chitosenuma** | **42.79** | **141.70** | **0.02** | **12** | **+/-** | **0.68** | **3** |
| **16** | **Konuma(Oshima)** | **41.97** | **140.66** | **3.77** | **12** | **+/-** | **0.14** | **2** |
| **17** | **Oonuma(Oshima)** | **42.00** | **140.69** | **5.30** | **12** | **+/-** | **0.14** | **1** |
| **18** | **Kokkuri** | **42.89** | **140.43** | **0.04** | **12** | **+/-** | **0.59** | **1** |
| **19** | **Hangetsu** | **42.85** | **140.76** | **0.05** | **12** | **+/-** | **0.49** | **1** |
| **20** | **Teshiopankenuma** | **45.03** | **141.72** | **3.54** | **12** | **+/-** | **0.04** | **N/A** |
| **21** | **Teshiopenkenuma** | **45.07** | **141.71** | **1.34** | **12** | **+/-** | **0.67** | **2** |
| **22** | **Poronuma** | **45.28** | **142.21** | **1.97** | **12** | **+/-** | **0.14** | **2** |
| **23** | **Kutcharo** | **45.14** | **142.33** | **8.13** | **6** | **+/-** | **0.08** | **N/A** |
| **24** | **Kabutonuma** | **45.22** | **141.69** | **0.80** | **12** | **+/-** | **0.64** | **2** |
| **25** | **Kushu** | **45.43** | **141.04** | **0.53** | **12** | **+/-** | **0.38** | **2** |
| **26** | **Chimikeppu** | **43.64** | **143.88** | **1.05** | **12** | **+/-** | **0.31** | **2** |
| **27** | **Toutsurutou** | **43.91** | **144.57** | **0.40** | **12** | **+/-** | **0.52** | **1** |
| **28** | **Saroma** | **44.14** | **143.82** | **151.36** | **6** | **+/-** | **0.00** | **N/A** |
| **29** | **Touya** | **42.60** | **140.86** | **70.59** | **12** | **+/-** | **0.12** | **2** |
| **30** | **Kuttara** | **42.50** | **141.18** | **4.68** | **12** | **+/-** | **0.19** | **3** |

| | | | | | | | | |
|---|---|---|---|---|---|---|---|---|
| 31 | Poroto | 42.56 | 141.36 | 0.34 | 12 | +/- | 0.18 | 2 |
| 32 | Shinonome | 43.27 | 143.14 | 0.05 | 12 | +/- | 0.39 | 1 |
| 33 | Shikaribetu | 43.28 | 143.12 | 3.60 | 12 | +/- | 0.18 | 1 |
| 34 | Horokayantounuma | 42.53 | 143.47 | 0.65 | 12 | +/- | 0.32 | 1 |
| 35 | Yuudounuma | 42.60 | 143.53 | 4.43 | 12 | +/- | 0.13 | 2 |
| 36 | Oikamanai | 42.55 | 143.49 | 1.49 | 12 | +/- | 0.22 | 2 |
| 37 | Choubushinuma | 42.66 | 143.61 | 1.22 | 12 | +/- | 0.48 | 1 |
| 38 | Takkobunuma | 43.11 | 144.48 | 1.31 | 12 | +/- | 0.73 | 2 |
| 39 | Akkeshi | 43.05 | 144.89 | 32.26 | 6 | +/- | 0.01 | N/A |
| 40 | Hichiripputo | 43.05 | 145.01 | 3.65 | 6 | +/- | 0.17 | N/A |
| 41 | Touro | 43.15 | 144.54 | 6.25 | 12 | +/- | 0.07 | N/A |
| 42 | Shirarutoro | 43.18 | 144.50 | 1.80 | 12 | +/- | 0.72 | 2 |
| 43 | Mashuu | 43.58 | 144.53 | 19.18 | 12 | +/- | 0.15 | 2 |
| 44 | Kussyaro | 43.63 | 144.33 | 79.39 | 12 | +/- | 0.06 | N/A |
| 45 | Akan | 43.45 | 144.10 | 13.23 | 12 | +/- | 0.11 | 1 |
| 46 | Tarou | 43.43 | 144.14 | 0.01 | 12 | + | 0.54 | N/A |
| 47 | Panketo | 43.48 | 144.18 | 2.85 | 12 | +/- | 0.16 | 3 |
| 48 | Rausu | 44.03 | 145.08 | 0.40 | 12 | +/- | 0.18 | 2 |
| 49 | Juusan | 41.03 | 140.36 | 18.05 | 6 | +/- | 0.02 | N/A |

| | | | | | | | | |
|---|---|---|---|---|---|---|---|---|
| **50** | **Towada** | **40.46** | **140.88** | **61.05** | **12** | **+/-** | **0.06** | **N/A** |
| **51** | **Anenuma** | **40.71** | **141.33** | **1.52** | **12** | **+/-** | **0.13** | **1** |
| **52** | **Ogawara** | **40.78** | **141.33** | **61.95** | **N/A** | **-** | **-0.01** | **N/A** |
| **53** | **Usorisan** | **41.32** | **141.09** | **2.67** | **12** | **+/-** | **0.22** | **1** |
| **54** | **Tappinuma** | **40.91** | **140.37** | **1.17** | **N/A** | **-** | **-0.05** | **N/A** |
| **55** | **Itobatakenoike** | **40.55** | **139.97** | **0.02** | **12** | **+/-** | **0.50** | **3** |
| **56** | **Nagaike** | **40.56** | **139.98** | **0.00** | **12** | **+** | **0.81** | **N/A** |
| **57** | **Hakkeinoike** | **40.56** | **139.96** | **0.01** | **12** | **+** | **0.65** | **N/A** |
| **58** | **Ouikehigashiike** | **40.56** | **139.97** | **0.04** | **12** | **+/-** | **0.40** | **3** |
| **59** | **Ooike-higashiike** | **40.54** | **139.97** | **0.09** | **12** | **+/-** | **0.43** | **3** |
| **60** | **Nigoriike** | **40.54** | **139.98** | **0.02** | **12** | **+** | **0.64** | **N/A** |
| **61** | **Tamokinuma** | **40.88** | **141.35** | **1.56** | **12** | **+/-** | **0.19** | **1** |
| **62** | **Ichiyanaginuma** | **40.91** | **141.36** | **1.69** | **N/A** | **+/-** | **0.04** | **N/A** |
| **63** | **Takahokonuma** | **40.94** | **141.33** | **5.49** | **12** | **+/-** | **0.23** | **3** |
| **64** | **Obuchinuma** | **40.96** | **141.35** | **3.50** | **12** | **+/-** | **0.19** | **3** |
| **65** | **Uchinuma** | **40.86** | **141.32** | **0.91** | **12** | **+/-** | **0.22** | **2** |
| **66** | **Onawashiro** | **39.85** | **140.98** | **0.02** | **12** | **+/-** | **0.64** | **1** |
| **67** | **Mangokuura** | **38.43** | **141.40** | **7.18** | **12** | **+/-** | **0.01** | **N/A** |
| **68** | **Izunuma** | **38.72** | **141.10** | **3.31** | **12** | **+/-** | **0.78** | **2** |

| | | | | | | | | |
|---|---|---|---|---|---|---|---|---|
| **69** | **Naganuma** | **38.70** | **141.13** | **3.06** | **12** | **+/-** | **0.76** | **2** |
| **70** | **Torinoumi** | **38.03** | **140.91** | **1.35** | **N/A** | **-** | **-0.03** | **N/A** |
| **71** | **Asanainuma** | **40.15** | **140.01** | **1.00** | **12** | **+/-** | **0.24** | **2** |
| **72** | **Hachirougata** | **39.92** | **140.01** | **27.65** | **12** | **+/-** | **0.03** | **N/A** |
| **73** | **Tazawa** | **39.73** | **140.66** | **25.72** | **N/A** | **+/-** | **0.06** | **N/A** |
| **74** | **Hakuryuu** | **38.06** | **140.18** | **0.06** | **12** | **+/-** | **0.52** | **1** |
| **75** | **Inawashiro** | **37.48** | **140.09** | **103.14** | **12** | **+/-** | **0.08** | **N/A** |
| **76** | **Matsukawaura** | **37.80** | **140.97** | **6.04** | **12** | **-** | **-0.09** | **N/A** |
| **77** | **Ozenuma** | **36.93** | **139.30** | **1.81** | **12** | **+/-** | **0.35** | **2** |
| **78** | **Akimoto** | **37.66** | **140.13** | **3.63** | **12** | **+/-** | **0.41** | **1** |
| **79** | **Onogawa** | **37.67** | **140.10** | **1.72** | **12** | **+/-** | **0.46** | **1** |
| **80** | **Sobara** | **37.69** | **140.07** | **0.31** | **12** | **+/-** | **0.40** | **1** |
| **81** | **Hibara** | **37.69** | **140.05** | **10.69** | **12** | **+/-** | **0.12** | **1** |
| **82** | **Sennba** | **36.37** | **140.46** | **0.29** | **12** | **+/-** | **0.06** | **N/A** |
| **83** | **Nishiura** | **36.04** | **140.37** | **167.60** | **12** | **-** | **-0.05** | **N/A** |
| **84** | **Ushikunuma** | **35.95** | **140.12** | **3.07** | **12** | **+/-** | **0.08** | **N/A** |
| **85** | **Sugaonuma** | **36.01** | **139.92** | **0.48** | **12** | **+** | **0.68** | **N/A** |
| **86** | **Kitaura** | **36.03** | **140.57** | **34.68** | **12** | **+/-** | **0.05** | **N/A** |
| **87** | **Sotonasakaura** | **35.91** | **140.60** | **5.76** | **12** | **-** | **-0.07** | **N/A** |

| | | | | | | | | |
|---|---|---|---|---|---|---|---|---|
| **88** | **Hinuma** | **36.28** | **140.50** | **9.37** | **12** | **-** | **0.00** | **N/A** |
| **89** | **Chuuzennji** | **36.74** | **139.46** | **11.80** | **12** | **+/-** | **0.02** | **N/A** |
| **90** | **Yuno** | **36.80** | **139.42** | **0.32** | **12** | **+/-** | **0.15** | **2** |
| **91** | **Haruna** | **36.48** | **138.87** | **1.23** | **6** | **+/-** | **0.15** | **N/A** |
| **92** | **Inbanuma** | **35.75** | **140.20** | **4.82** | **12** | **+/-** | **0.42** | **1** |
| **93** | **Teganuma** | **35.86** | **140.03** | **3.69** | **12** | **+/-** | **0.25** | **3** |
| **94** | **Shinsei** | **35.36** | **139.21** | **0.01** | **12** | **+** | **0.73** | **N/A** |
| **95** | **Ashino** | **35.21** | **139.01** | **6.89** | **6** | **+/-** | **0.08** | **N/A** |
| **96** | **Toyanogata** | **37.89** | **139.05** | **1.35** | **12** | **+/-** | **0.07** | **N/A** |
| **97** | **Hyou** | **37.84** | **139.24** | **0.12** | **12** | **+/-** | **0.66** | **2** |
| **98** | **Kamo** | **38.07** | **138.43** | **4.81** | **N/A** | **+/-** | **0.05** | **N/A** |
| **99** | **Kahokugata** | **36.65** | **136.67** | **4.29** | **12** | **-** | **-0.02** | **N/A** |
| **100** | **Kibagata** | **36.37** | **136.45** | **1.09** | **12** | **-** | **-0.05** | **N/A** |
| **101** | **Kitagata** | **36.27** | **136.24** | **2.14** | **12** | **+/-** | **0.06** | **N/A** |
| **102** | **Shibayamagata** | **36.35** | **136.37** | **1.97** | **12** | **-** | **-0.02** | **N/A** |
| **103** | **Ouchigata** | **36.92** | **136.81** | **0.62** | **12** | **+/-** | **0.06** | **N/A** |
| **104** | **Hiruga** | **35.61** | **135.89** | **0.92** | **12** | **+/-** | **0.05** | **N/A** |
| **105** | **Kugushi** | **35.60** | **135.91** | **1.42** | **6** | **+/-** | **0.01** | **N/A** |
| **106** | **Suigetsu** | **35.59** | **135.88** | **4.20** | **N/A** | **+/-** | **0.01** | **N/A** |

| | | | | | | | | |
|---|---|---|---|---|---|---|---|---|
| **107** | **Suga** | **35.58** | **135.90** | **0.92** | **N/A** | **+/-** | **0.05** | **N/A** |
| **108** | **Mikata** | **35.57** | **135.88** | **3.58** | **12** | **+/-** | **0.21** | **2** |
| **109** | **Motosu** | **35.46** | **138.59** | **4.70** | **12** | **+/-** | **0.17** | **2** |
| **110** | **Shouji** | **35.49** | **138.61** | **0.46** | **12** | **+** | **0.19** | **N/A** |
| **111** | **Yamanaka** | **35.42** | **138.87** | **6.40** | **12** | **+/-** | **0.01** | **N/A** |
| **112** | **Sai** | **35.50** | **138.69** | **2.13** | **12** | **+/-** | **0.15** | **2** |
| **113** | **Kawaguchi** | **35.51** | **138.75** | **5.70** | **N/A** | **+** | **0.14** | **N/A** |
| **114** | **Suwa** | **36.05** | **138.08** | **12.90** | **12** | **+/-** | **0.20** | **2** |
| **115** | **Kizaki** | **36.56** | **137.84** | **1.66** | **12** | **+/-** | **0.20** | **2** |
| **116** | **Nakatsuna** | **36.60** | **137.84** | **0.12** | **12** | **+/-** | **0.42** | **3** |
| **117** | **Aoki** | **36.61** | **137.85** | **1.73** | **12** | **+** | **0.16** | **N/A** |
| **118** | **Ina** | **36.05** | **138.46** | **0.10** | **12** | **+/-** | **0.31** | **3** |
| **119** | **Hijiri** | **36.49** | **138.07** | **0.07** | **12** | **+/-** | **0.24** | **3** |
| **120** | **Nojiri** | **36.83** | **138.22** | **4.43** | **12** | **+/-** | **0.19** | **3** |
| **121** | **Sanaru** | **34.71** | **137.69** | **1.17** | **12** | **+/-** | **0.08** | **N/A** |
| **122** | **Hamana** | **34.74** | **137.59** | **64.51** | **N/A** | **-** | **-0.02** | **N/A** |
| **123** | **Ippeki** | **34.93** | **139.11** | **0.19** | **N/A** | **+** | **0.22** | **N/A** |
| **124** | **Inohana** | **34.79** | **137.56** | **5.42** | **N/A** | **-** | **-0.01** | **N/A** |
| **125** | **Shiraishi** | **34.11** | **136.24** | **0.46** | **12** | **+** | **0.15** | **N/A** |

| | | | | | | | | |
|---|---|---|---|---|---|---|---|---|
| **126** | **Biwa** | **35.29** | **136.09** | **669.83** | **N/A** | **-** | **-0.05** | **N/A** |
| **127** | **Nishino** | **35.16** | **136.12** | **2.13** | **12** | **+/-** | **0.12** | **1** |
| **128** | **Hira** | **35.05** | **135.92** | **0.13** | **12** | **+/-** | **0.13** | **2** |
| **129** | **Matsunokinai** | **35.31** | **136.05** | **0.13** | **12** | **+** | **0.47** | **N/A** |
| **130** | **Omatsunai** | **35.23** | **135.96** | **0.08** | **12** | **+** | **0.27** | **N/A** |
| **131** | **Yogo** | **35.52** | **136.19** | **1.75** | **12** | **+/-** | **0.20** | **2** |
| **132** | **Asokai** | **35.57** | **135.18** | **4.79** | **N/A** | **+/-** | **0.03** | **N/A** |
| **133** | **Hanare** | **35.69** | **135.04** | **0.37** | **12** | **+/-** | **0.16** | **1** |
| **134** | **Kumihama** | **35.63** | **134.91** | **7.17** | **N/A** | **+/-** | **0.02** | **N/A** |
| **135** | **Koyamaike** | **35.51** | **134.15** | **6.99** | **12** | **-** | **0.00** | **N/A** |
| **136** | **Nakanoumi** | **35.48** | **133.20** | **87.03** | **6** | **+/-** | **0.02** | **N/A** |
| **137** | **Tougouike** | **35.48** | **133.89** | **4.09** | **N/A** | **-** | **-0.02** | **N/A** |
| **138** | **Shinji** | **35.45** | **132.96** | **78.99** | **12** | **-** | **-0.03** | **N/A** |
| **139** | **Jinzai** | **35.33** | **132.68** | **1.15** | **N/A** | **-** | **0.00** | **N/A** |
| **140** | **Banryuu** | **34.68** | **131.81** | **0.09** | **N/A** | **+** | **0.49** | **N/A** |
| **141** | **Oomi** | **34.41** | **131.18** | **0.23** | **12** | **+** | **0.64** | **N/A** |
| **142** | **Ezu** | **32.77** | **130.75** | **0.33** | **12** | **+** | **0.43** | **N/A** |
| **143** | **Kamiezu** | **32.78** | **130.74** | **0.15** | **12** | **+** | **0.57** | **N/A** |
| **144** | **Shidaka** | **33.26** | **131.46** | **0.08** | **12** | **+/-** | **0.13** | **3** |

| | | | | | | | | |
|---|---|---|---|---|---|---|---|---|
| **145** | **Ikeda** | **31.24** | **130.56** | **10.92** | **N/A** | **+/-** | **0.09** | **N/A** |
| **146** | **Namakoike** | **31.87** | **129.87** | **0.52** | **12** | **+** | **0.24** | **N/A** |
| **147** | **Unagiike** | **31.22** | **130.61** | **1.20** | **N/A** | **+/-** | **0.08** | **N/A** |
| **148** | **Satsuma** | **31.51** | **130.34** | **0.11** | **12** | **+** | **0.41** | **N/A** |

**Table S2.** Leaf thickness, fresh weight, chlorophyll concentration, and reflectance at red (670 nm) and NIR (800 nm) of leaf samples. n=5 (sd).

| Species | Thickness | FW | [Chl] | | Red | NIR |
|---|---|---|---|---|---|---|
| | mm | $g/m^2$ | $mg/m^2$ | μg/gFW | % | % |
| *Lemna minor* | 0.18 (0.01) | 191 (28) | 124 (19) | 648 (16) | 5.1 (1.6) | 37.3 (3.5) |
| *Azolla filiculoides* | 0.67 (0.03) | 410 (72) | 267 (45) | 653 (53) | 1.0 (0.2) | 34.8 (1.2) |
| *Salvinia molesta* | 0.74 (0.18) | 232 (55) | 280 (18) | 1271 (331) | 2.4 (0.4) | 42.1 (3.0) |
| *Limnobium Laevigatum* | 5.35 (0.70) | 878 (71) | 295 (60) | 336 (68) | 6.0 (0.7) | 50.6 (3.2) |
| *Eichhornia crassipes* | 0.43 (0.06) | 292 (36) | 439 (102) | 1553 (555) | 10.3 (3.0) | 57.4 (3.9) |
| *Nymphaea spp* | 0.73 (0.16) | 333 (39) | 498 (71) | 1518 (288) | 14.4 (4.4) | 54.9 (6.8) |
| *Trapa japonica* | 0.38 (0.02) | 250 (7) | 300 (38) | 1200 (148) | 26.1 (2.5) | 70.6 (4.0) |
| *Arabidopsis thaliana* | 0.26 (0.02) | 258 (16) | 355 (40) | 1388 (237) | 2.8 (0.6) | 35.4 (1.7) |
| *Ligustrum lucidum* | 0.32 (0.02) | 298 (21) | 473 (72) | 1593 (285) | 12.9 (1.7) | 64.1 (2.2) |
| *Verbascum thapsiforme* | 0.74 (0.12) | 235 (13) | 644 (125) | 2729 (393) | 7.0 (0.3) | 59.8 (1.6) |
| *Kalanchoe CLONEKOE* | 2.6 (0.50) | 2301 (371) | 297 (43) | 133 (41) | 18.7 (2.2) | 49.2 (4.0) |
